# The transcriptome landscape of developing barley seeds reveals H3K27me3 dynamics in endosperm tissues

**DOI:** 10.1101/2023.07.26.550659

**Authors:** Martin Kovacik, Anna Nowicka, Jana Zwyrtková, Beáta Strejčková, Isaia Vardanega, Eddi Esteban, Asher Pasha, Kateřina Kaduchová, Maryna Krautsova, Jan Šafář, Nicholas J. Provart, Rüdiger Simon, Ales Pecinka

## Abstract

Cereal grains are an important food and feed. To provide comprehensive spatiotemporal information about biological processes in developing seeds of cultivated barley, we performed a transcriptomic study of the embryo, endosperm, and seed maternal tissues collected from 4 to 32 days after pollination. Weighted gene co-expression network and motif enrichment analyses pointed out specific groups of genes and transcription factors with possible impacts on regulating barley seed tissue development. We defined a set of tissue-specific marker genes and families of transcription factors for functional studies of the pathways controlling barley grain development. Assessment of selected groups of chromatin regulators revealed that epigenetic processes are highly dynamic and likely to play a major role during barley endosperm development. Repressive modification H3K27me3 is globally reduced in endosperm tissues and at specific developmental and storage compound genes. Altogether, this atlas uncovers the complexity of the developmental regulation of gene expression in barley grains.

**Teaser:** Spatiotemporal profiling of developing barley seeds revealed loss of H3K27me3 and changes in gene expression in endosperm.

## Introduction

Seeds are a crucial stage in the life cycle of many plants allowing the survival of long periods of unfavorable conditions and colonization of new sites. High nutritive value and long storability made seeds one of the prime targets of plant breeding. Among seeds, cereal grains are one of the most valuable agronomic products (*1*). Cultivated barley (*Hordeum vulgare* L. subsp. *vulgare*) is the fourth most important cereal worldwide and is used for feed (70%), malt (21%), and direct consumption (9%) (*2*). Barley is an important diploid temperate cereal model species with a long history of research and plentiful genetic resources (*3*). Its grains are composed of three main compartments: diploid embryo combining one maternal and one paternal genome, triploid endosperm harboring two maternal genomes and one paternal genome, and seed maternal tissues (SMTs) of maternal origin (*4*). The developing embryo undergoes a series of differentiation steps. It consists of the embryonic axis (coleoptile, plumule, shoot apical meristem, radicle, coleorhiza) and the scutellum at maturity. Endosperm development involves an initial stage where nuclei divide without cell division, creating a coenocyte, which accumulates simple sugars. Subsequently, the nuclei become separated by cell walls during cellularization, and endosperm further differentiates into specific domains (*5*). The mature endosperm is composed of four main cell types, including the central starchy endosperm (CSE) that serves as the main storage of complex sugars, the aleurone layer (AL), embryo-surrounding region (ESR), and basal endosperm transfer layer (BETL) (*6*). Finally, an outer layer of SMTs, consisting of a pericarp and two layers of seed coats, maintains stable conditions and protects the embryo and endosperm. The molecular mechanisms governing the development of individual seed parts of tissues remain only poorly understood in barley.

Spatial and temporal analysis of gene expression is an effective approach for the basic characterization of developmental programs. Previously, several studies explored transcriptional profiles of developing barley grains and their parts. Analysis of expressed sequence tags revealed transcriptional reprogramming in young seed maternal and filial tissues (0 to 7 days after pollination; DAP) and early to late whole caryopses (*7*). The seed maternal tissues showed up-regulation of genes encoding many protein and lipid activating enzymes and down-regulation of reactive oxygen species (ROS)-scavenging enzyme genes. This suggested the mobilization of storage compounds and programmed cell death (PCD). The filial tissues contained highly expressed genes involved in cell growth, including cell wall biosynthesis. The first whole genome barley grain expression studies were conducted on Affymetrix Barley1 GeneChips with samples from dissected embryos and endosperm tissues 16 to 25 DAP and whole caryopses 5 to 16 DAP (*8, 9*). These studies described the dynamics in the expression of genes encoding components of many key metabolic and hormonal pathways during grain development and their potential roles during germination. RNA sequencing of whole caryopses helped to explore the role of RNA editing in barley grain development (*10*). The most recent transcriptomic study provided a spatially resolved cellular map for germinating barley seeds (*11*).

Here, we performed a comprehensive transcriptome profiling on barley seed tissues at different stages of development with an aim to create an atlas of gene expression and thus provide detailed information on the temporal and spatial distribution of the key molecular processes acting during barley grain development. The focus on the nucleus-driven processes will be fundamental for future functional studies of the role of the epigenome in barley grain development.

## Results

### Generating tissue-specific transcriptome profiles of developing barley seeds

To identify transcriptional signatures of the major tissues of developing barley seeds, we performed RNA sequencing (RNA-seq) of the dissected embryo, endosperm, and SMTs of the cultivar (cv.) Morex at 4, 8, 16, 24, and 32 DAP (Fig. 1, A and B). Sample hierarchical clustering revealed a strict grouping of the tissues and time points, except for the 4 DAP endosperm that clustered separately, indicating its unique transcriptome (Fig. 1C and fig. S1). This pattern was further corroborated by principal component analysis (PCA), which revealed a predominant clustering according to DAP (PC1) and tissues (PC2), explaining in total 70% of the variability (Fig. 1D). The 4 and 8 DAP endosperm samples were notably distant from the later time points, indicating a massive transcriptional reprogramming during endosperm proliferation and cellularization. The closer distance between the 24 and 32 DAP samples for both embryo and endosperm suggested minor transcriptional changes at the end of grain development. No major changes in clustering were observed by inspecting PC3 (18% of the variability; fig. S2, A and B). To view the seed transcriptome in the context of the whole barley plant, we performed PCA together with the transcriptomes of eight different tissues (*12*). The PC1 and PC2 explained 46% and 21% of the variance, and the overall distribution was defined by the seed samples. The vegetative tissues consisting of a root, shoot, nodule, and inflorescences clustered together with the germinating embryo and were positioned close to the 8 DAP embryo and 4 DAP SMTs. The caryopsis 5 and 15 days after anthesis grouped with the 4 and 16 DAP endosperm, respectively (Fig. 1E and fig. S2, C and D), suggesting that the endosperm dominates the caryopsis transcriptome profile.

**Fig. 1.**
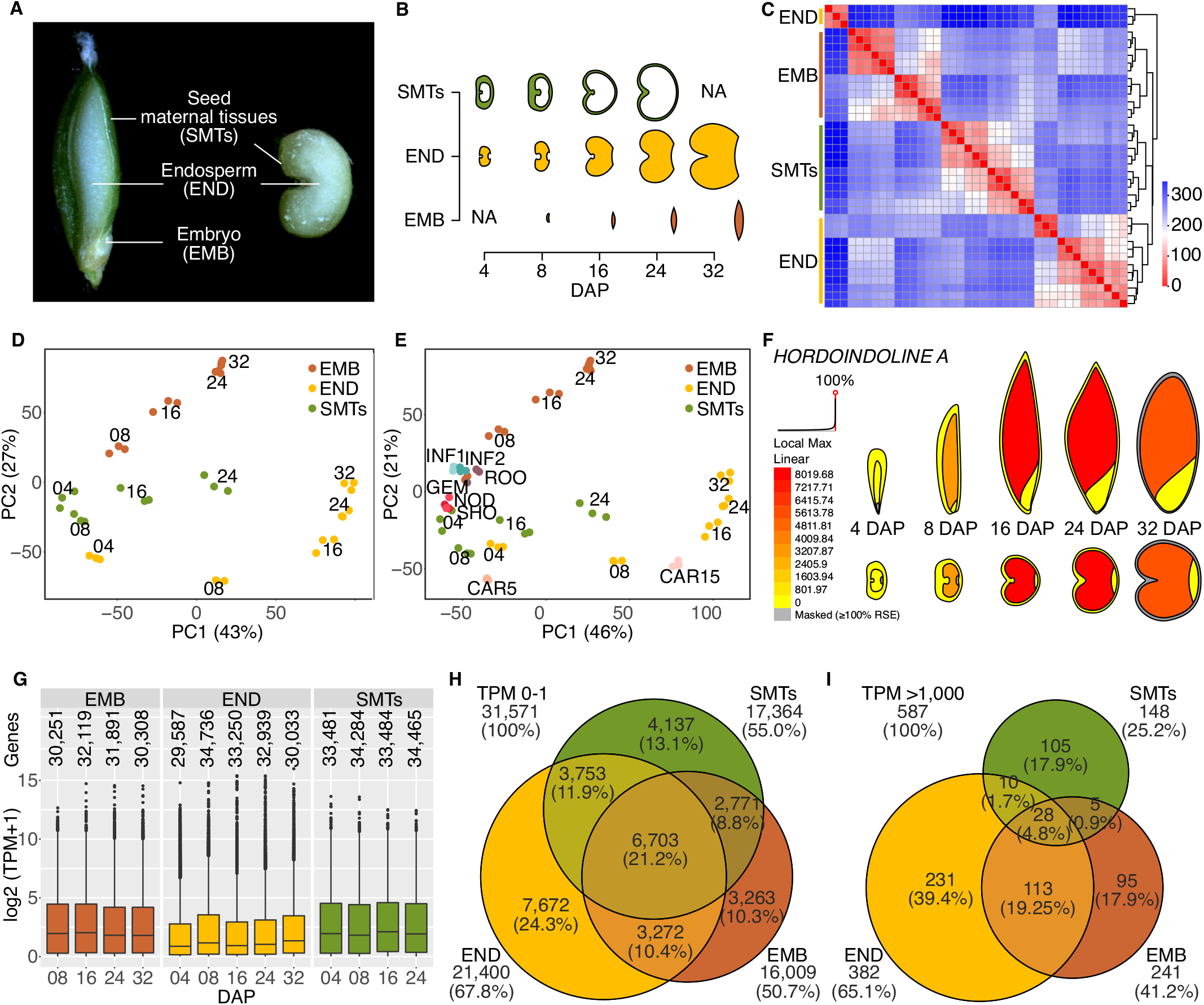
Transcriptomes of developing barley grain. (**A-B**) Overview of the analyzed tissues and time points used for transcriptomic analysis. DAP -Days after pollination; NA - not analyzed. (**C**) Sample distance heatmap of 39 seed-specific tissue transcriptomes hierarchically clustered based on their similarity. The scale shows the sample distance. (**D**) The variance of the 39 samples represented by principal component (PC) analysis for the embryo (orange), endosperm (yellow), and seed maternal tissues (green). The numbers within the graph indicate DAP and three close spots represent biological triplicates. The number next to PC indicates variance. (**E**) Principal component analysis of seed tissues used in (C) in combination with other barley tissues (*12*). ROO – root, GEM – germinating embryo, NOD – nodule, SHO – shoot, INF1 and INF2 – developing inflorescence of 5- and 10-mm length, CAR5 and CAR15 – caryopsis 5 and 15 DAP. (**F**) Visualization of interactive heatmap at ePlant Barley. Heatmap in transverse (upper) and sagittal (lower) sections shows TPM expression levels of *HORDOINDOLINE A* in different tissues of developing barley grain. (**G**) Boxplots of expression for genes with non-zero expression (source values in data S1). (**H**) Venn diagram showing numbers of lowly expressed genes (0-1 TPM during entire grain development). (**I**) Venn diagram of the highest expressed genes (>1,000 TPM).

Comparison of expression levels for individual genes (shown as transcripts per million, TPM; data S1) revealed striking differences between the tissues. To allow easy visualization of the data in a user-friendly format, we integrated our dataset into Barley ePlant browser on The Bio-Analytic Resource for Plant Biology (BAR) platform (Fig. 1F; https://bar.utoronto.ca/eplant_barley/). The endosperm median expression was 0.96 – 1.35 TPM compared to 1.80 – 2.03 TPM in the embryo and 1.81 – 2.11 TPM in SMTs (Fig. 1G). Genes with low expression (TPM 0 – 1; n = 31,571) included 67.8% endosperm (n = 21,400), 50.7% embryo (n = 16,009), and 55.0% SMT (n = 17,364) expressed genes (Fig. 1H). Hence, endosperm has a 2-fold lower median expression compared to the other tissues and the highest number of very weakly expressed genes. Surprisingly, endosperm also contained the majority of the genes with the highest expression (> 1000 TPM; endosperm n = 382; embryo n = 241; SMTs n = 148; Fig. 1I). Despite this, the proportion of endosperm genes was constantly decreasing for expression from 1 to 1000 TPM (fig. S3, A, B and C). The genes with the highest expression in endosperm were significantly enriched in negative regulation of proteolysis, defense response, endosperm development, response to wounding, lipid transport, and cell wall macromolecule catabolic process (table S1), representing some of the key processes known to play a role at different stages of endosperm development.

### Grain development is associated with extensive transcriptional changes

To assess the tissue and stage-specific gene expression, we plotted the differentially expressed genes (DEGs) at one or more time points for each tissue (Fig. 2A). During embryo development, the major transcriptional changes occurred from 8 to 24 DAP with 8,952 (8 to 16 DAP) and 10,340 (16 to 24 DAP) DEGs, respectively (FDR-adjusted P < 0.05; Fig. 2A, fig. S4A and data S2). On the contrary, the transition from 24 to 32 DAP was accompanied by a lower number of 4,959 DEGs (FDR-adjusted P < 0.05). In endosperm, there were massive and unique transcriptional changes from 4 to 8 DAP, as indicated by the 15,990 DEGs with 7,584 up-regulated (5,181 unique) and 8,406 down-regulated (4,688 unique) cases (FDR-adjusted P < 0.05; Fig. 2A, fig. S4B and data S2). The number of endosperm DEGs gradually decreased toward 32 DAP. There was a similar trend to embryo also in SMTs, with more DEGs between the early and middle time points (5,604 between 4 and 8 DAP and 8,264 between 8 and 16 DAP; FDR-adjusted P < 0.05; Fig. 2A, fig. S4C and data S2) and less at the late stage of development (3,953 DEGs at 16 to 24 DAP).

**Fig. 2.**
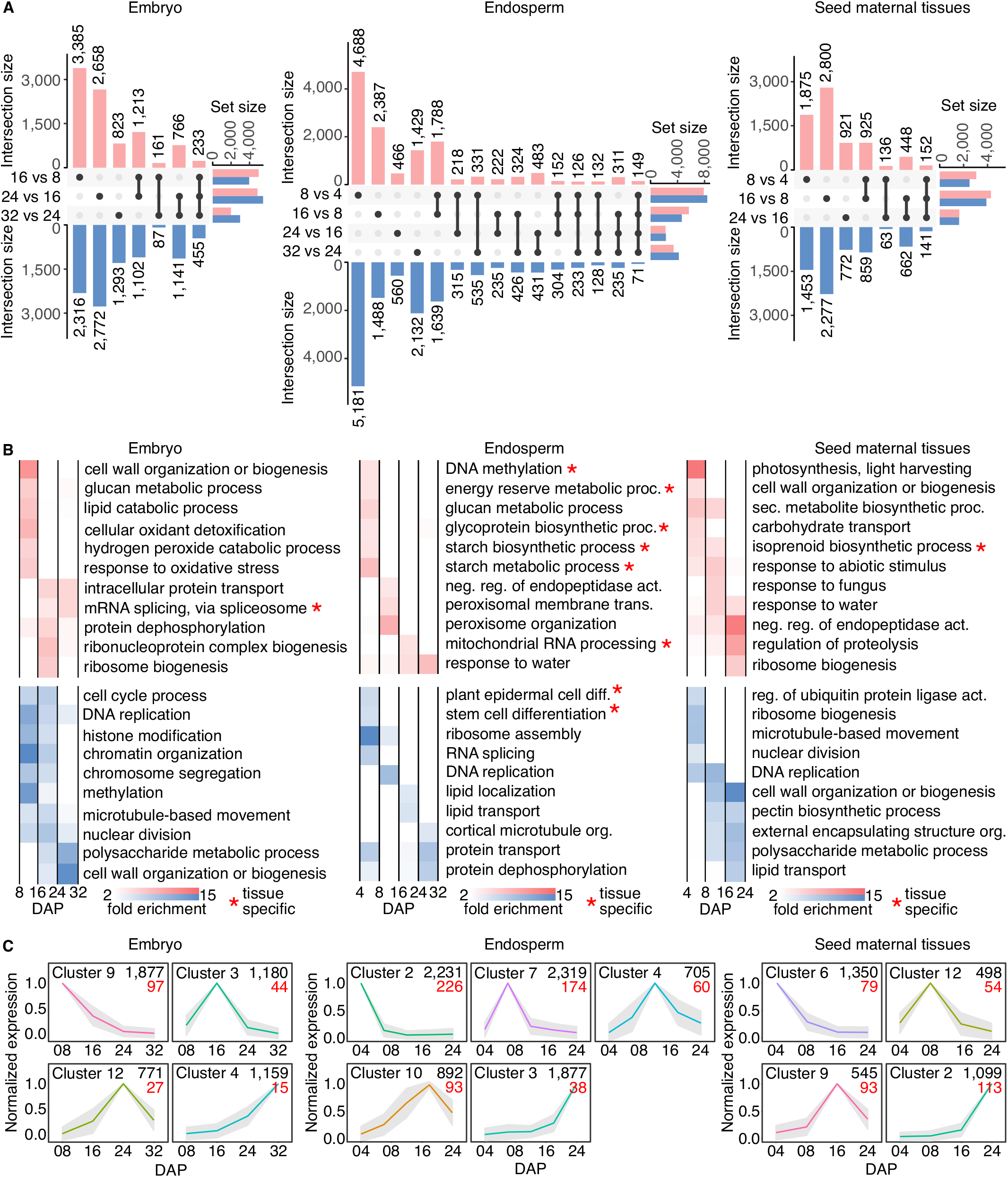
Tissue and timepoint specificity of gene expression during barley grain development. (**A**) The UpSet plots show up-regulated (red) and down-regulated (blue) differentially expressed genes (DEGs) in embryo, endosperm, and seed maternal tissues. The set size on the x-axis defines the total number of DEGs between two subsequent experimental time points. The y-axis shows the number of DEGs (intersection size) in single stages (single black dots) and their combinations (connected dots; source data in data S2). (**B**) Gene ontology (GO) term enrichment of DEGs (FDR-adjusted P < 0.05) between developmental timepoints in embryo, endosperm, and seed maternal tissues. The top representative GO categories among up-regulated (red) or down-regulated (blue) genes are shown. Color saturation corresponds to fold enrichment (full list in tables S2 and S3). Tissue-specific GO categories are indicated by a red asterisk. (**C)** K-means co-expression clusters of DEGs, peaking in single consecutive developmental time points in embryo, endosperm, and seed maternal tissues. The black numbers in the upper right corner indicate gene count in individual clusters. The red numbers indicate subset of tissue specific marker genes. DAP – days after pollination (the full list in fig. S5 and data S3).

We used the sets of DEGs to associate transitions between individual DAPs with over- or under-representation of Gene Ontology (GO)-based biological processes. The 8 DAP embryo development was associated with cell division (GOs: cell cycle, DNA replication, histone, and chromatin modification) and its reduction before 16 DAP (Fig. 2B and table S2). A 16 DAP embryo transcriptome indicated intense ribosome biogenesis and cell wall synthesis. However, these GOs are typical for many actively growing tissues and were also shared to a large extent with endosperm and SMTs (table S2 and S3). The embryo-specific GO term was mRNA splicing via spliceosome enriched after 16 DAP and even more after 24 DAP (Fig. 2B). This is consistent with the accumulation of mRNAs in developing seeds and their translation during germination (*13*). In endosperm, there were enriched many tissue-specific processes linked with storage compounds (GOs: lipid-, starch-, glucan-, glycoprotein-biosynthesis and metabolism; Fig. 2B, table S2 and table S3), which underlies the role of endosperm as the main nutritious tissue in grains. Furthermore, there was a tissue-specific expression of genes coding for DNA methylation factors from 4 to 8 DAP in the endosperm. The most prominent GO terms in SMTs included: photosynthesis and cell wall development (both peaking at 8 DAP), and a wave of expression from the fungi and abiotic stress-responsive factors (Fig 2B, table S2 and table S3). The unique GO term of SMTs was the upregulation of isoprenoid biosynthesis from 4 to 16 DAP. Accumulation of isoprenoids such as hormones (brassinosteroids, gibberellins, abscisic acid) has an important impact on seed nutritional and physiological quality, i.e. seed longevity and dormancy (*14*).

We used K-means clustering of DEGs to define the specific genes that could serve as molecular markers for individual tissues and DAPs (Fig. 2C, fig. S5 and data S3). The embryo (n = 17,214), endosperm (n = 21,889), and SMTs (n = 15,034) DEGs were divided into 12, 13, and 14 co-expression clusters, respectively, where four to five clusters showed expression peaking at single consecutive experimental points. Each cluster was searched for tissue-specific genes defined as having less than 5% TPM in other seed tissues (Fig. 2C and fig. S5 – red numbers). The most of the stage and tissue specific genes in embryo and endosperm were found in 4 and 8 DAP specific clusters.

### Specific promoter motifs are associated with transcriptional regulation of seed development

To identify correlation patterns of mRNA levels among genes across our data we performed weighted gene co-expression network analysis (WGCNA) for each tissue. The identified modules were organized temporally as to their profile being characteristic for the early, middle, or late stages of development (Fig. 3A, fig. S6 and data S4). Notably, the majority of the genes accumulated at early (EMB_M2, n = 1,883; END_M2/M7/M6, n = 1,962; and SMT_M1, n = 1,572) and late (EMB_M1, n = 1,934; END_M1/M2, n = 1,838; and SMT_M3, n = 2,497) modules in individual tissues. This suggests that early and late seed developmental phases are more dynamic concerning transcriptional reprogramming.

**Fig. 3.**
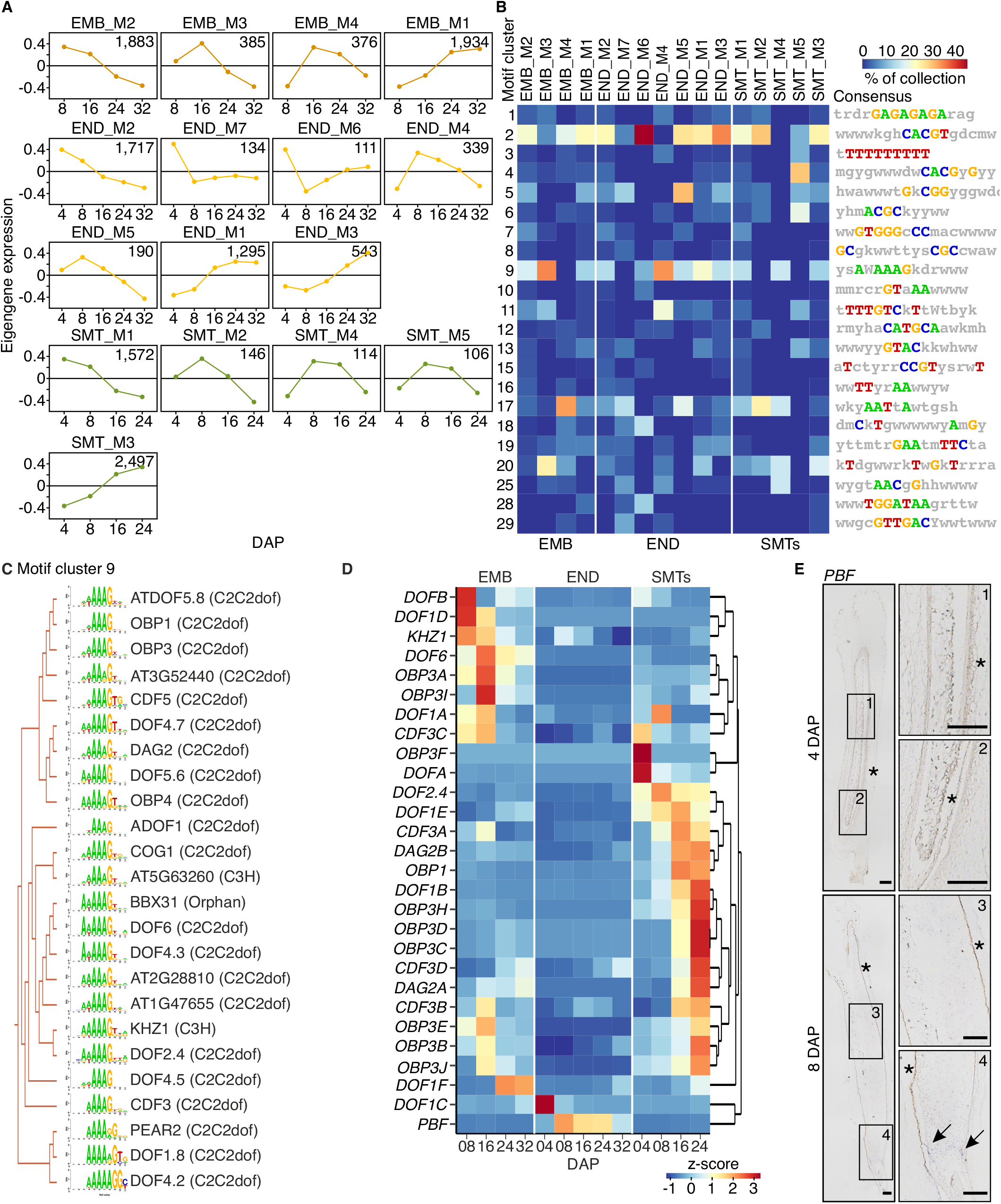
Gene co-expression network analysis and promoter motif enrichment. (**A**) Display of selected weighted gene co-expression network analysis (WGCNA) modules (the full list of modules is provided in data S4 and fig. S6). The graphs show eigengene expression in each module. The numbers in the upper right corner indicate gene count in individual clusters. EMB – embryo, END – endosperm, SMTs – seed maternal tissues, DAP – days after pollination. (**B**) Analysis of transcription factor (TF) binding motifs in individual WGCNA modules shown in (A) by screening promoters of barley genes against the Arabidopsis TF binding motif database and clustering. The consensual motif in each motif cluster (MC, rows) is shown on the right side and its representation (%) in the WGCNA module is indicated by the color scale. The heatmap depicts some motifs from individual clusters identified in each WGCNA module containing at least 10 motifs (full list in table S5). (**C**) Hierarchical clustering of TF binding sequence motifs in MC2 with the *in silico*-predicted binding Arabidopsis TFs and their families (in parentheses). (**D**) Heatmap of hierarchically clustered expression for barley orthologs of Arabidopsis TFs from (C) in EMB, END, and SMTs (source data in table S7). (**E**) RNA *in situ* hybridization of *PBF* gene in barley grains at 4 and 8 DAP. Arrows indicate signal deposition in the endosperm. Scale bars 200 m. Asterisks indicate native staining of the testa (not a signal).

To obtain information about transcription factors (TFs) and regulatory motifs important during barley grain development, we performed promoter motif enrichment analysis. The sequences -1,500 to -1 bp from the transcription start site of genes included in WGCNA modules, representing their presumable cis-regulatory regions, were analyzed for the presence of known TF binding motifs using the *Arabidopsis thaliana* HOMER database (*15*). This resulted in collections of significantly enriched motifs (p-val < 0.05; data S5). The motifs were grouped based on their similarity into clusters (table S4 and table S5), and the proportion of major motif clusters (MCs) in each collection was investigated (Fig. 3B). The MC2 and MC9 were enriched across many WGCNA modules of all tissues and their consensual motifs corresponded to G-box (CACGT) and P-box (AWAAG), respectively (Fig. 3B). Next, we investigated individual MC2 DNA binding motifs in detail (fig. S7A). The commonly included TF families were basic/helix-loop-helix (bHLH) and basic region/leucine zipper (bZIP) known to bind the G-box, and TKACGT motif variants (fig. S7A). To test whether the predicted TFs might be further supported by our data, we identified barley orthologs of Arabidopsis TFs (table S6), and investigated their expression (fig. S7B and table S6). The expression profile of many of these TFs appeared early or late during grain development, e.g. barley homolog of *ABA INSENSITIVE 5* (*ABI5*) showed a late expression profile in embryo and endosperm. In Arabidopsis, *ABI5* is involved in signaling during seed maturation and regulates a subset of *LATE EMBRYOGENESIS ABUNDANT* (*LEA*) genes (*16*). Many *LEA* genes were highly expressed in the late embryo module, and some also in the late endosperm module (data S4). The MC9 was enriched particularly at middle embryo and endosperm development and contained the characteristic Dof TF family binding P-box motif (A/T)AAAG (Fig. 3, B and C and table S7). Dof TFs regulate seed storage protein synthesis in maize and barley (*17, 18*). Furthermore, a barley Dof TF *PROLAMIN BINDING FACTOR* (*PBF*) activates the expression of storage compounds in endosperm by binding to the P-box motif present in the promoters of many storage protein genes (*18*). Our data confirmed spatial and temporal expression patterns of this gene in barley endosperm (Fig. 3, D and E). As the P-box motif was enriched in the middle endosperm module, we investigated the most expressed genes in the END_M4 module and observed many enzymes involved in storage protein synthesis, such as sucrose synthase, alpha-glucan-branching enzymes, and starch synthase as well as low molecular weight glutenin subunit (data S4).

### The barley endosperm differentiation program is initiated before cellularization

The fully differentiated cereal endosperm contains at least four main morphologically distinguishable domains (*19*). Although markers for individual endosperm domains were identified across cereals, their number is limited for barley (*20*–*25*). Therefore, we performed a comparative analysis of 12 markers for individual endosperm domains described in maize and rice. By reciprocal BLAST, we identified 29 homologs in barley (table S8). The candidate markers were expressed specifically in the endosperm. Interestingly, they typically reached a maximum expression at younger (4 or 8 DAP) stages (Fig. 4, fig. S8 and table S8), suggesting a biased selection. Nevertheless, several genes could be potentially good late endosperm development markers in barley.

**Fig. 4.**
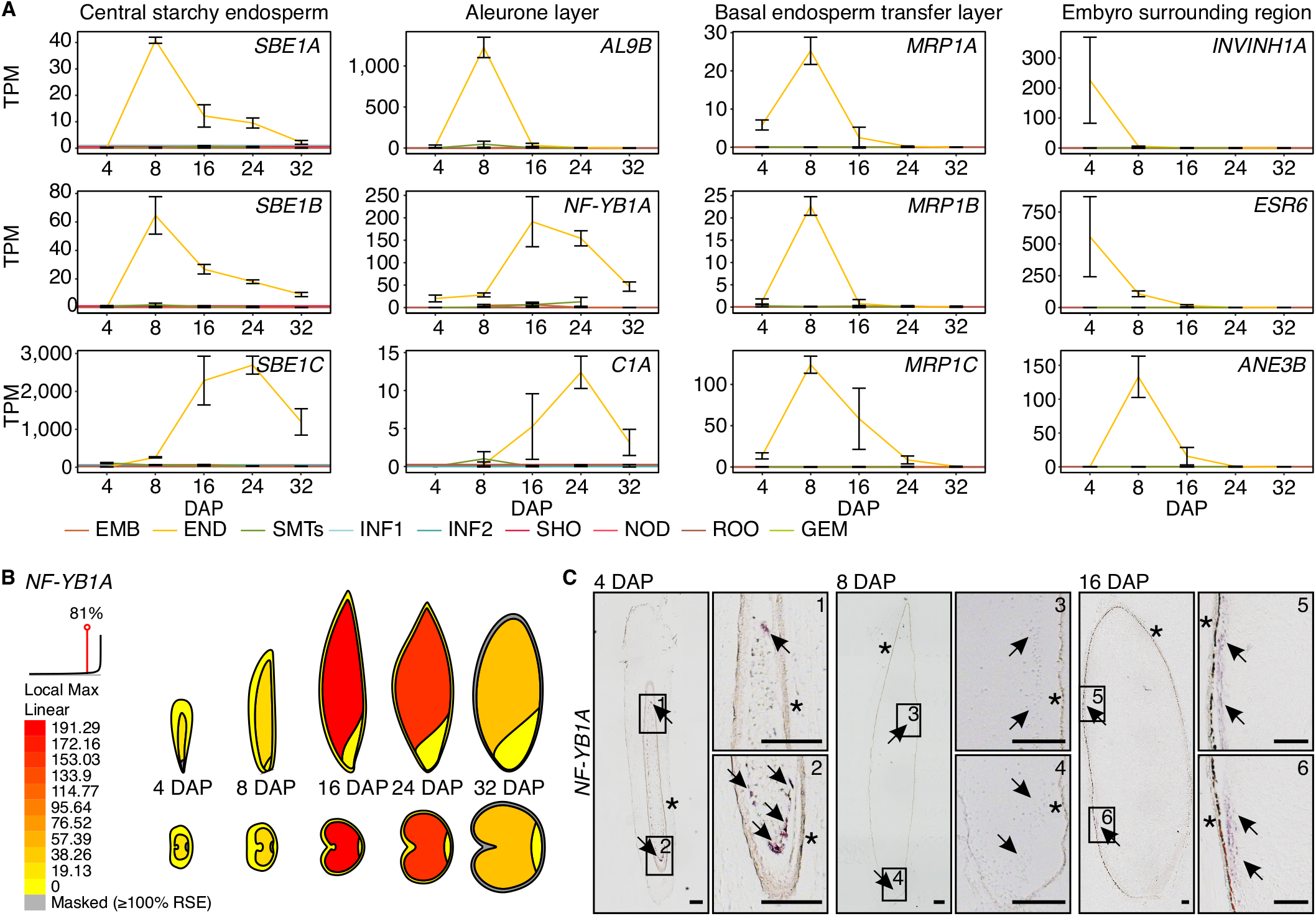
Expression from markers of endosperm domains in barley. (**A**) Expression profiles of barley orthologs of selected endosperm marker genes in other cereals grouped according to the domain of expression - central starchy endosperm, aleurone layer, basal endosperm transfer layer, and embryo surrounding region across different barley tissues (*12*). EMB – embryo, END – endosperm, SMTs – seed maternal tissues, ROO – root, GEM – germinating embryo, NOD – nodule, SHO – shoot, INF1 and INF2 – developing inflorescence 5 and 10 mm, CAR5 and CAR15 – caryopsis 5 and 15 days after pollination (DAP). Error bars indicate standard deviation (the full list in table S8 and fig. S8). (**B**) RNA *in situ* hybridization of *NF-YB1A* gene in barley grains at 4, 8, and 16 days after pollination (DAP). Arrows indicate signal deposition in the aleurone region. The asterisk shows a naturally colored testa layer that does not represent RNA *in situ* signal. Scale bars 200 m. (**C**) Visualization *NF-YB1A* expression (transcripts per million) in different tissues of developing barley grain at Barley ePlant.

The timing of starch granule accumulation differs between cereals. It starts soon after cellularization around 6 DAP in barley, whereas it begins at round 10 DAP in maize (*26, 27*). The expression of the three barley copies of CSE marker *STARCH BRANCHING ENZYME 1* (*SBE1A-C*) was firstly detected at 8 DAP, but only the *SBE1A* increased its expression until it peaked at 24 DAP (Fig. 4A). The maximal expression of barley homologs of maize *ALEURONE9* (*AL9A* and *AL9B*), rice *NUCLEAR TRANSCRIPTION FACTOR Y SUBUNIT B-1A* (*NF-YB1A*) and maize *COLORED ALEURONE1* (*C1A*) were observed at 8, 16 and 24 DAP, respectively, indicating that AL is established later during endosperm development (Fig. 4, A and B and fig. S8). Indeed, we confirmed the expression of *NF-YB1A* at 16 DAP using RNA *in situ* hybridization (Fig. 4C). Findings in maize suggest that AL differentiates from the outer layers of endosperm cells between 6 and 10 DAP soon after alveolation and the first periclinal division of the cellularized endosperm (*6*). As a control, we also analyzed 4 DAP endosperm. Surprisingly, *NF-YB1A* was also expressed there and showed an interesting accumulation of the signal at the basal pole of the seed (Fig. 4C). This suggested that the aleurone identity is defined in some endosperm nuclei already before cellularization and starts at the embryonic pole. Similarly, BETL markers *EMBRYO SAC/BASAL ENDOSPERM TRANSFER LAYER/EMBRYO SURROUNDING REGION 1* paralogs A and D (*EBE1A-D*), resembling antimicrobial proteins and *MYB-RELATED PROTEIN 1* paralogs *A* to *C* (*MRP1A-C*), which promote expression of two unrelated transfer cell–specific genes in maize, initiated their expression at 4 DAP and peaked at 8 DAP (Fig. 4A and fig. S8). Transfer cell fate specification in barley apparently occurs during a narrow temporal window of coenocyte development, similarly to maize (*28*). The ESR markers *INVERTASE INHIBITOR 1A* (*INVINH1A*) likely involved in sugar management, defensin-type gene *EMBRYO SURROUNDING REGION 6A* (*ESR6A*), and two genes with similarity to antifungal proteins *ANDROGENIC EMBRYO 1* paralogs *A* to *B* (*ANE1A-B*) and *ANDROGENIC EMBRYO 3* paralogs *A* to *B* (*ANE3A-B*), reached their peak of expression at 4 DAP (Fig. 4A and fig. S8). This corresponds to findings in maize, where ESR cells differentiate upon completion of the endosperm cellularization phase (*29, 30*). Hence, we confirmed the specificity of known endosperm markers from other cereals in barley; however, their expression was in some cases shifted. The AL, BETL, and ESR markers were already expressed at 4 and 8 DAPs, suggesting that the specificity of certain endosperm domains is already defined at their founder nuclei before the cellularization.

### Multiple pathways are controlled by H3K27me3 in barley endosperm

The GO term enrichment analyses indicated a role of specific chromatin-based pathways during barley grain development (Fig. 2B, table S2 and table S3). To explore this direction, we identified barley orthologs of two selected groups previously connected with seed development - histones and the subunits of the POLYCOMB REPRESSIVE COMPLEX 2 (PRC2) (table S9).

We identified 188 barley histone genes corresponding to all canonical forms and common plant variants (fig. S9 and table S10). In total, 152 out of 175 expressed histones (87%, TPM ≥ 1) were part of the endosperm k-means CL2 (including 30 copies of H3; 31 copies of H2A.W; and one H2A.Z), with a predominant expression during the coenocyte stage of endosperm development (Fig. 2C and data S3). The peak of expression in CL2 coincided with the period of DNA replication and nuclei division during coenocyte endosperm development. After cellularization, the expression of histone genes decreased, and only a few, mostly paralogous copies of non-canonical variants remained expressed (H2A.W, H3.3; fig. S9 and table S10). The initial stages of embryo and SMTs were also marked by the peak of histone expression, but the overall strength of expression was lower than in endosperm (fig. S9). There was a prominent expression from the recently described seed-specific histone H2B.S variant (*31*) in 16 DAP and later embryo stages. The H2B.S was silent in endosperm and SMTs. This indicates dynamic epigenetic control and change in nucleosome composition during endosperm and embryo development.

The PRC2 complex installs tri-methylation of lysine 27 at histone H3 (H3K27me3). This modification is known to transcriptionally repress its target genes and thus contribute to developmental transitions, including endosperm cellularization (*32*). Arabidopsis PRC2 consists of four evolutionary conserved subunits FERTILIZATION INDEPENDENT ENDOSPERM (FIE), MULTICOPY SUPPRESSOR OF IRA 1 (MSI1), EMBRYONIC FLOWER 2 (EMF2), and the catalytic subunit represented by two homologs SET DOMAIN GROUP 1 and 10 (SDG1, a.k.a. CURLY LEAF, CLF and SDG10, a.k.a. SWINGER, SWN). All PRC2 subunits are present in cereals (*33*). We found that the barley genome contains single copies of *FIE* and *SDG10/SWN* and multiple copies of *MSI1, EMF2*, and *SDG1/CLF*. At least one copy of each PRC2 subunit was well expressed (TPM > 10) throughout the whole embryo and SMT development (Fig. 5A and table S11). The pattern in the endosperm was more complex. Genes coding for *EMF2A*/*B, SDG1A*/*B*, and *SDG10* were not, or only weakly, expressed at 4 DAP but there was an expression from *FIE* and *MSI1A*/*B*. The latter could be also due to the dual role of MSI1 in PRC2 and histone H3.1 chaperone complex Chromatin Assembly Factor-1 (CAF-1) (*34*). This suggests no or very little function of the PRC2 in the early (4 DAP) endosperm development in barley. From 8 DAP onwards, there was an expression from *EMF2A, FIE, MSI1B*, and *SDG10*, while both *SDG1* copies remained not or weakly expressed. The high degree of expression from *SDG10* versus *SDG1* indirectly suggests that SDG10/SWN might be a functional homolog of the Arabidopsis endosperm-specific PRC2 catalytic subunit SDG5/MEDEA in barley. To estimate whether this expression pattern might have any phenotypic effects, we performed H3K27me3 immunostaining on isolated nuclei with different DNA contents (C-values) from 8 DAP and 24 DAP endosperm and 24 DAP embryo (Fig. 5B). There was an intense H3K27me3 signal at the telomeric pole in embryo nuclei of all ploidies and DAPs. Such signal distribution is caused by the accumulation of H3K27me3 at the ends of Rabl-organized barley chromosomes (*35, 36*). On the contrary, the H3K27me3 signal was weaker at 8 DAP endosperm nuclei and almost absent at 24 DAP (Fig. 5B and fig. S10), suggesting a cytologically detectable global loss of H3K27me3 from high C-value and older endosperm nuclei. Speculatively, SDG10/SWN containing PRC2 might have a narrower range of genomic targets that could be linked to certain functions, such as genomic imprinting.

**Fig. 5.**
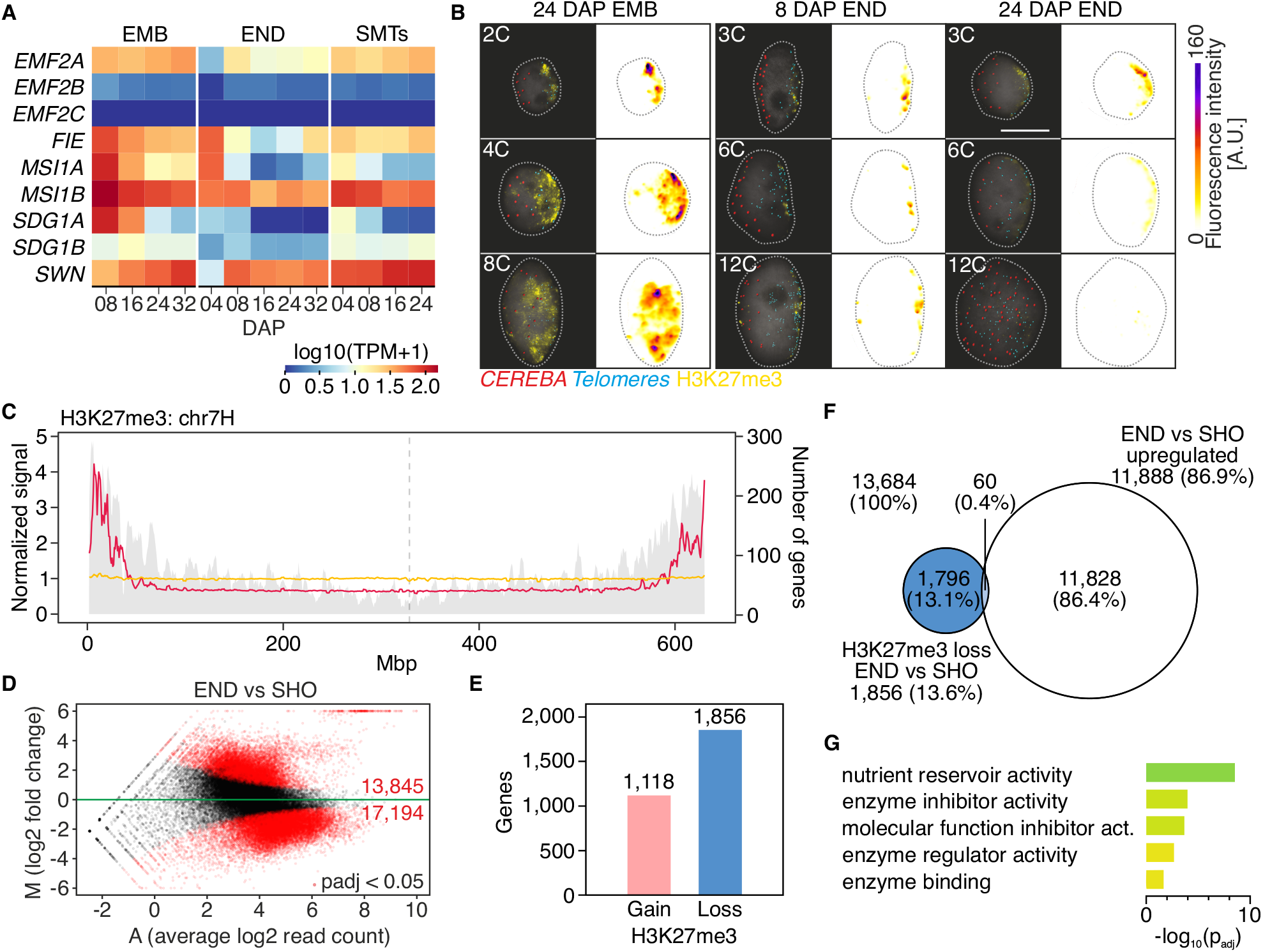
Dynamics of expression from Polycomb repressive complex 2 (PRC2) genes in barley grain tissues. (**A**) Heatmap of expression from genes coding PRC2 complex subunits in the embryo (EMB), endosperm (END), and seed maternal tissues (SMTs) at different days after pollination (DAP, source data in table S11). (**B**) Black background images show representative embryo and endosperm nuclei of different C-values collected at 8 and 24 DAP. DNA was stained with DAPI (grey), H3K27me3 was immunostained (yellow), and *CEREBA* centromeric (red) and telomeric (blue) repeats were visualized by fluorescence *in situ* hybridization and signal segmentation. White background images show H3K27me3 immunostaining fluorescence signal intensities in arbitrary units (A.U.). Scale bar = 10 μm. Raw images and 3D image segmentation pictures of the nucleus surface, and immunostaining and FISH signals are presented in fig. S10. (**C**) Normalized signal abundance of H3K27me3 in endosperm (yellow) and 10-cm whole seedling (*36*) (magenta). The gray background signal is gene density (secondary y-axis; full list is provided in fig. S11) across chromosome 7H. (**D**) MAplot showing genomic intervals with differential signal intensities between endosperm and 10-cm seedling (SHO). Intervals passing the threshold (Benjamini-Hochberg FDR-adjusted p-value < 0.05) are in red. The red numbers indicate number of genomic intervals with significant gain (M > 0) or loss of H3K27me3 (M < 0). Source data are provided in data S6. (**E**) Number of genes corresponding to genomic regions with significant gain and loss of H3K27me3 (full list provided in data S7). (**F**) Venn diagram showing the number of genes with loss of H3K27me3 in 10-cm seedlings (SHO; based on ChIP-seq data from Baker et al., 2015) and genes up-regulated at 8, 16, 24, or 32 DAP endosperm. (**G**) Gene ontology (GO) term enrichment of genes with loss of H3K27me3 and significant up-regulation in endosperm (source data in table S13).

To investigate H3K27me3 levels in endosperm at greater resolution, we performed chromatin immunoprecipitation and sequencing (ChIP)-seq on 16 DAP endosperm samples, and compared the signals to H3K27me3 ChIP-seq data from seedlings (*36*) (Fig. 5C, fig. S11 and fig. S12A). As expected, the H3K27me3 signals concentrated on gene-rich chromosome termini in seedlings but were strongly reduced in endosperm. Although the overall trend was biased towards H3K27me3 reduction in endosperm, the changes were more complex when looking on individual H3K27me3 peaks (fig. S12B, C and D). The total number of H3K27me3 peaks was higher in endosperm, nevertheless they were shorter and smaller compared to seedlings. We focused on the most prominent peaks (fold enrichment ≥ 5) and performed differential analysis. We found 17,194 regions with a significant H3K27me3 loss and 13,845 regions with gain (log2FoldChange < 0 or > 0, respectively, Benjamini-Hochberg FDR-adjusted p-value < 0.05) in endosperm relative to shoot (Fig. 5D and data S6). The genomic regions within loss category included 1,856 and those from gain region 1,118 genes (data S7). To assess direct role of H3K27me3 in their transcriptional regulation, we searched which of these genes were significantly up-regulated or down-regulated (padj < 0.05) in at least one analyzed endosperm time point (Fig. 5F, fig. S12E, table S12 and table S14).

Among the 60 (0.4%) endosperm H3K27me3-depleted and up-regulated genes were several storage compound genes including *LOW-MOLECULAR-WEIGHT GLUTENIN SUBUNITs, C-HORDEIN* and *OMEGA SECALIN* (table S12). Interestingly, other such genes included inhibitors of sugar, and protein degradation (*SERPIN*) and were expressed in a seed stage-dependent manner (table S12). The *INVERTASE INHIBITORs* that block hydrolysis of sucrose to fructose or glucose were highly expressed at 8 DAP but not during subsequent stages, possibly allowing feeding of the growing embryo or endosperm. The inhibitor of protein degradation *SERPIN* was expressed from 16 DAP (table S12). This suggests that the accumulation of storage carbohydrates and proteins is accompanied by simultaneous inhibition of their degradation in the endosperm tissues in H3K27me3 dependent manner, which was supported also by the enrichment of related GO terms (Fig. 5G and table S13).

There were 238 (1.4%) H3K27me3-enriched significantly down-regulated genes from at least one analyzed endosperm time point (fig. S12E, table S14). This cluster was dominated by two prominent groups. The first were putative *SENESCENCE-ASSOCIATED PROTEINs* (n = 72) which might be regulating tissue maturation by inhibiting specific proteases (*37*). The second cluster included 27 copies of the RNA Pol II subunit *MEDIATOR OF RNA POLYMERASE II TRANSCRIPTION SUBUNIT 12* (*MED12*) (table S14). MED12 could be linked with transcriptional regulation of specific genes in barley. In Arabidopsis, it contributes to regulation of flowering genes and the mutants show late-flowering phenotype (*38*). Among enriched GO categories we observed terms related to respiration and generation of energy (fig. S12F and table S15).

This shows an interesting dynamic of chromatin-controlled processes in barley endosperm. Expression of histone genes appears critical during DNA replication and nucleus division at the coenocyte stage. During grain filling and senescence, several biological pathways appear to be controlled by the H3K27me3 modification via PRC2.

### Genomic imprinting of specific genes is conserved among cereals

Studies in Arabidopsis showed that H3K27me3 plays an important role in the epigenetic regulation of uniparental gene expression by genomic imprinting in early endosperm development (*39*). Until now, only eight imprinted genes were identified in barley based on the homology with imprinted genes in rice and subsequent validation (*40*). We took an analogous approach with a broader dataset of 155, 156, and 697 imprinted genes from rice, maize, and bread wheat, respectively (*41*–*45*), and performed a comparative search for evolutionarily conserved imprinted genes in barley endosperm tissues. We identified 449 barley orthologs (Fig. 6A and table S16) with 21 (4.3%) cases shared by at least two species. Such a low number indicates a relatively low evolutionary conservation of imprinted genes across the main grass species. We checked the expression of the 21 conserved candidates in our seed transcriptomic data and identified four main expression pattern groups (Fig. 6B and table S17). Group 1 contained nine genes expressed across all seed tissues. Group 2 had four genes expressed in embryo and SMTs but mostly lowly expressed or silent in the endosperm. Group 3 consisted of seven genes with expression restricted to endosperm. Lastly, group 4 contained a single gene expressed at early endosperm, 8 DAP SMTs, and silent in the embryo.

**Fig. 6.**
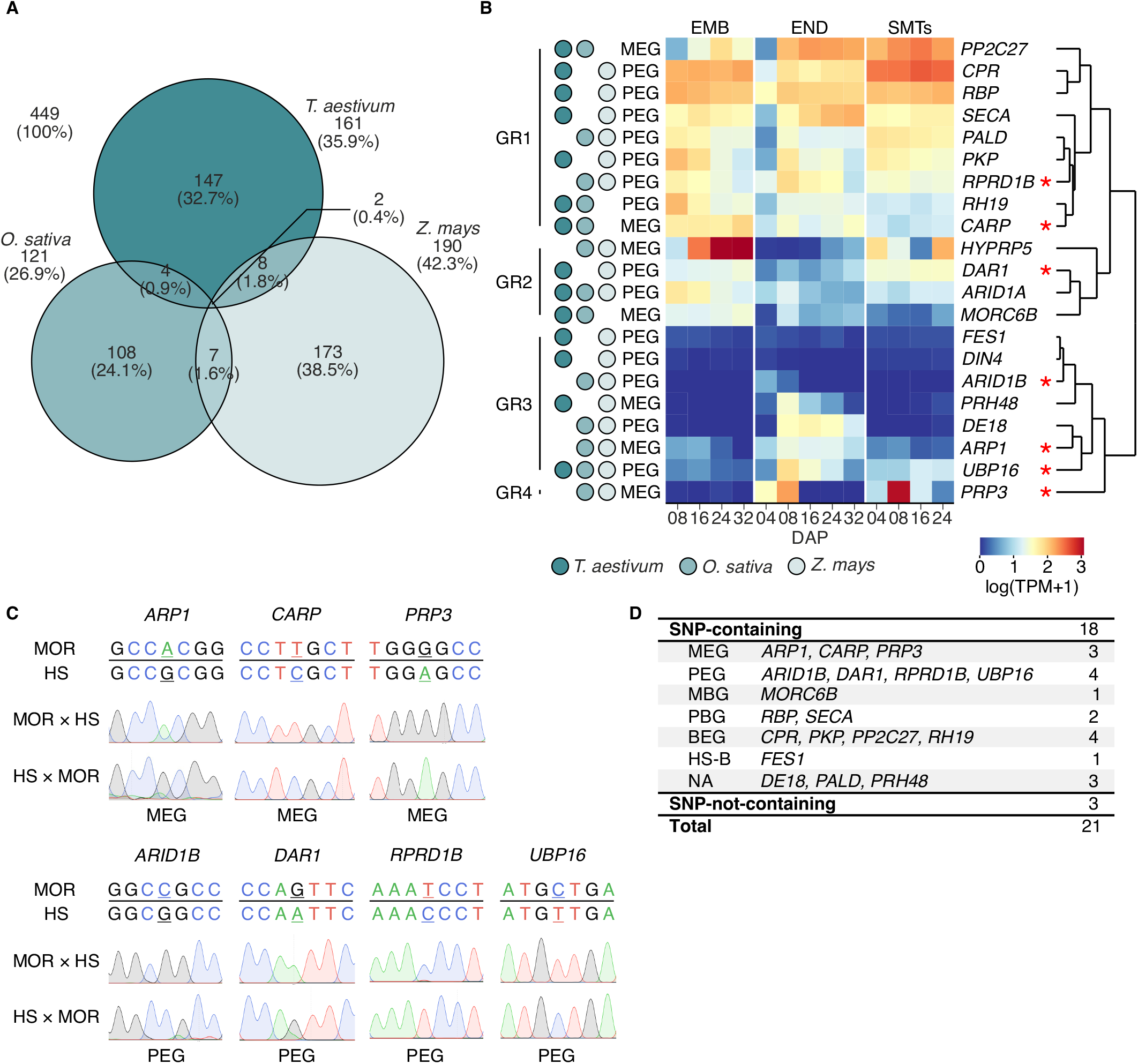
Identification and validation of imprinted genes in barley endosperm. (**A**) Venn diagram showing the number of genes imprinted in single and multiple cereal species (source data in table S16). (**B**) Heatmap of expression of barley orthologs of 21 genes imprinted in multiple cereal species in the embryo (EMB), endosperm (END), and seed maternal tissues (SMTs) at different days after pollination (DAP). The heatmap is partitioned into groups based on different expression profiles in seed tissues. MEG – maternally expressed gene, PEG – paternally expressed gene, * - imprinted in barley (source data in table S17) (**C**) Validation of SNPs for selected imprinted genes in 8 DAP endosperm tissues by Sanger sequencing. F1 reciprocal hybrids were obtained by crossing genetically distant cultivars Morex (MOR) and wild barley (HS). The maternal genotype is mentioned first in the crosses. The informative SNPs are underlined. (**D**) Summary of validation of imprinted genes in barley. MBG – maternally biased gene, PBG – paternally biased gene, BEG – biallelic expressed gene, HS-B – wild barley biased, NA – information not available.

To provide experimental validation, we tested whether the 21 candidates represent true imprinted genes in barley. Two genetically distant strains cv. Morex and *H. vulgare* subsp. *spontaneum* accession HOR 12560 (HS) were reciprocally crossed, and subjected to the allelic expression analysis of the candidate genes from 8 DAP endosperm. In total, 18 genes contained diagnostic SNPs and were tested for the parent-of-origin specific expression (Fig. 6C, D and fig. S12, S13). Amplification of cDNA of three genes *DEFECTIVE ENDOSPERM* (*DE18*), *PALADIN* (*PALD*) and *PROTEIN PHOSPHATASE HOMOLOG 48* (*PRH48*) was not successful and therefore these candidates remain unclassified. The *FRIGIDA-ESSENTIAL 1* (*FES1*) gene showed a HS genotype-specific expression. Four genes *CYTOCHROME P450 REDUCTASE* (*CPR*), *PROTEIN KINASE FAMILY PROTEIN* (*PKP*), *PROBABLE PROTEIN PHOSPHATASE 2C 27* (*PP2C27*) from group 1 were expressed from both parents. Three genes were tested as potentially imprinted because they showed either maternally (*MORC6B*) or paternally (*RBP* and *SECA*) biased expression. Seven of the total 18 tested genes (38.9%) were confirmed as imprinted in barley (Fig. 6D). Three maternally expressed genes (MEGs) included *CA-RESPONSIVE PROTEIN* (*CARP*), *RNA-BINDING PROTEIN* (*ARP1*) and *PROLINE-RICH PROTEIN* (*PRP3*). Wheat and rice MEG *CARP* was part of barley Group 1 and was expressed only weakly in endosperm compared to moderate expression in embryo. *ARP1* and *PRP3* were both MEGs in rice and maize and their expression was relatively specific to barley endosperm. An interesting spike of *PRP3* expression occurred in 8 DAP SMTs. We also confirmed four paternally expressed genes (PEGs) *REGULATION OF NUCLEAR PRE-MRNA DOMAIN-CONTAINING PROTEIN 1B* (*RPRD1B*), *DA1-RELATED 1* (*DAR1*), *AT-RICH INTERACTION DOMAIN 1B* (*ARID1B*) and *UBIQUITIN-SPECIFIC PROTEASE 16* (*UBP16*).

The *RPRD1B* and *ARID1B* genes were found as PEGs in rice and maize. *RPRD1Bs* are epsin N-terminal homology (ENTH) domain-containing proteins with roles in endocytosis and cytoskeletal regulation (*46*). Arabidopsis ARID1 is a transcriptional activator expressed in pollen development, which could be consistent with its role as PEG in barley. *DAR1* was previously found as PEG in wheat and maize (Fig. 6B). It is related to Arabidopsis *DA1*, an ubiquitin-activated peptidase which plays a role in the regulation of endoreduplication, determination of plant architecture and possibly maternal control of seed mass (*47, 48*). DA1 functions antagonistically with its direct substrate UBIQUITIN-SPECIFIC PROTEASE 15 (UBP15) in Arabidopsis (*49*). Interestingly, we confirmed another member of UBC family, *UBIQUITIN-SPECIFIC PROTEASE 16* (*UBP16*) as evolutionary conserved PEG in maize, rice, wheat (*41, 43*–*45*) and barley.

Finding seven imprinted genes encourages future genome-wide search in barley and provides an initial set of evolutionarily conserved candidates for the functional studies in barley.

## Discussion

Cultivated barley is a temperate crop of growing importance as a model species for cereal research (*3*). Here, we provided so far the most detailed atlas of gene expression during seed development in barley using the current version of genome assembly and annotation. The stage- and tissue-specific marker genes defined in our study serve as a solid basis for the follow-up biological investigations on the functional validation of crucial players involved in barley grain development. Although we paid most attention to endosperm tissues, the dataset offers equal resolution also for embryo and often neglected seed maternal tissues that play a key role in defining grain parameters, e.g. its size (*50*). The role of chromatin in governing seed development is studied in detail in the Arabidopsis model system and has revealed a number of epigenetic processes and molecular factors playing important roles (*51*). How much of this is conserved in cereals, including barley, remains largely unknown. We performed several pilot experiments that defined expression patterns of histones and PRC2 silencing complex. Both the dynamic expression changes and cytologically detectable reduction in H3K27me3 modification point to the pivotal role of chromatin in barley grain development and stimulate the need for further functional studies. Finally, we used a comparative approach to identify the first imprinted genes in barley that are conserved with other cereals. Several confirmed candidates point to the role of ubiquitin-based regulation of genomic imprinting in barley and grasses. Future whole genome studies will be needed to uncover the full spectrum of barley imprinted genes. In conclusion, our study will promote and facilitate further functional research on barley grain development and associated pathways.

## Materials and Methods

### Plant materials and growth conditions

Six-rowed spring barley *Hordeum vulgare* subsp. *vulgare* cv. Morex was used throughout the study. For analysis of impritend genes, also wild barley *Hordeum vulgare* subsp. *spontaneum* strain HOR 12560 (HS) was used. For plant growth, the seeds were germinated on moisturized cellulose tissue paper covered with one layer of filter paper for 3 days at 25°C in the dark. Germinating grains were moved into 5 × 5 cm peat pots containing a mixture of soil and sand (3/1; v/v) and were grown in a climatic chamber under a long-day regime (16 h day 20°C, 8 h night 16°C; light intensity 200 μmol m^-2^ s^-1^; humidity 60%). After 10 days, the plants were transferred into 15 × 15 cm pots containing the same soil and cultivated in the same conditions until flowering. The number of DAPs was defined by determining the day of self-pollination as described (*52*). The developing seeds were collected at 4, 8, 16, 24, and 32 DAPs in at least three biological replicates. Only the central row seeds from the middle of the spikelets were used. The 4 DAP embryo was omitted due to its small size. For sampling 8 DAP embryos, at least 10 embryos were pooled per biological replicate. For embryo samples at later stages, five embryos were pooled per biological replicate. Young (4 and 8 DAP) and older endosperm (>8 DAP) tissues were isolated from twenty or three seeds, respectively, per biological replicate as described (*52*). SMTs were isolated from at least five grains per biological replicate, irrespective of the stage of development. SMTs at 32 DAP were excluded due to their dry nature. Dissected tissues were immediately frozen in liquid nitrogen and stored at -80°C until use.

### RNA extraction, sequencing, and analysis

Total RNA was isolated using an RNeasy^®^ Plant Mini Kit (QIAGEN) and Spectrum™ Plant Total RNA Kit (Sigma-Aldrich) according to the manufacturer’s instructions including on-column DNase I (Sigma-Aldrich) treatment. The RNA quality was checked using Bioanalyzer 2100 with RNA 6000 Pico Chips (all Agilent). Samples with RNA integrity number >6.8 were processed into RNA-Seq mRNA libraries using NEBNext^®^ Ultra™ RNA Library Prep Kit for Illumina^®^ with poly-A selection. The mRNA-enriched libraries were sequenced as single-end 100 bp RNA seq reads on a HiSeq2500 instrument (Illumina). The raw reads were trimmed for adaptors by Trim Galore v.0.4.1 (*53*) and aligned to the *H. vulgare* cv. Morex reference genome v3 (*54*) using HiSat2 v.2.1.0 genomic mapper (*55*). Aligned reads were assigned to features and meta-features using Subread v.1.5.2 software (*56*) according to the genome annotation. Differential expression analysis was performed using DESeq2 v.1.24.0 package (*57*) in R v.3.6.3 software (*58*). A gene was declared to be significantly differentially expressed if the Benjamini-Hochberg FRD-adjusted p-value was <0.05. Published barley RNA-Seq data sets were retrieved at the NCBI SRA from Bioproject PRJEB3149 and analyzed as described above. The PCAs were done after applying the variance stabilization transformation (*59*). Venn diagrams were drawn using the eulerr v.6.1.0 package in R (*60*).

### Clustering analyses

For k-means clustering, unique DEGs from all tested combinations within the tissue were clustered using the k-means algorithm in R (*58*). K-means clustering was performed using the Hartigan-Wong algorithm with 1,000 iterations. An optimal number of clusters was determined by statistical testing using a gap statistics method (*61*). For WGCNA a gene co-expression network was constructed for each tissue with the raw read counts after the *rlog* transformation using the WGCNA library in R (*62, 63*). An adjacency matrix was made using the soft thresholded Pearson correlation (power = 18) among 5,000 genes with the most varying expression among experimental points. Hierarchical clustering was performed for grouping the genes with highly similar co-expression patterns. The modules were identified using the Dynamic Hybrid Tree cut algorithm (*64*) keeping the minimum size of the module to 15 and DeepSplit set to 4 to produce fine clusters. Each module was represented by color coding, with 12-15 modules detected depending on the tissue. The expression profile of each module was summarized by module eigengene defined as its first principal component. The probes that did not fit any of the main modules were placed into the “unspecific” module that was removed from further analysis.

### A seed view in Barley ePlant

The Barley ePlant framework (*65*) was modified to accept V3 Barley gene identifiers (*54*). To create a new Seed view in the Barley ePlant, the data described in this publication in TPM were databased on the Bio-Analytic Resource for Plant Biology’s server (*66*). An SVG image depicting the parts of the seed that were sampled in this work was generated and an XML file linking specific parts of the image with database sample names was manually created to configure this new view.

### GO term enrichment and annotation of transcription factors

GO annotation was used to perform an enrichment analysis by the topGO v.2.44.0 package in R (*67*). Redundant GO terms were filtered using the web-based tool REVIGO (*68*) with default settings and general terms were filtered using size selection (*69*). Terms with a size sufficient for robust statistical analysis (n > 100) and fold enrichment > 3 were investigated. Transcription factors were classified into families (data S8) based on the presence of specific domains according to PlantTFDB (*70*).

### Cis-motif identification and clustering

For cis-motif identification and enrichment analysis, 1500 bp upstream sequences from the predicted start codon (ATG) of all WGCNA module genes were used. The analysis was carried out using the “findMotif.pl” program from the HOMER suite (*15*) that performs known motif identification and enrichment analysis with default parameters. The enrichment of identified motifs was calculated respective to all 1500 bp background sequences. Collections of identified motifs in each WGCNA module were post-filtered for plant motifs and clustered using the RSAT (*71*) with default parameters.

### RNA *in situ* hybridization

Barley seeds were harvested at 4, 8, and 16 DAP, fixed with 4% freshly prepared paraformaldehyde (w/v), with 2% Tween-20 and 2% Triton X-100 (pH 7, adjusted by HCl) for 1 h under vacuum. For increased fixation efficiency vacuum was broken every 10 min and applied again. Subsequently, seeds were transferred into fresh fixatives and stored overnight at 4°C, dehydrated using ethanol series, cleared by ROTIHistol series, and embedded into Paraplast. Tissue sections of 10 μm were cut with a Reichert-Jung 2030 microtome and attached to Adhesion Slides Superfrost Ultra Plus (Thermo Fisher Scientific). DNA templates for synthesis of RNA probes were amplified from cDNA (reverse transcribed by RevertAid H Minus First Strand cDNA Synthesis Kit; Thermo Fisher Scientific) by PCR. Sense and antisense digoxigenin (DIG)-labeled RNA probes were amplified using gene-specific primers containing T7 promoter sequences (table S18) and DIG-UTP (Thermo Fisher Scientific) according to the manufacturer’s instructions. After purification, the efficiency of DIG labeling was verified by a modified dot blotting hybridization (*72*). For hybridization, slides with tissue sections were washed in ROTIHistol, rehydrated, and treated with 0.2 M HCl for 10 min, pronase (0.125 mg mL^-1^) for 10 min, 0.2% glycine for 2 min, 4% formaldehyde for 10 min and acetic anhydride (1% in 0.1 M Triethanolamine pH 8.0). Hybridizations with denatured probes were carried out at 50°C using the hybridization buffer containing 100 μL 10× salts, 400 μL deionized formamide, 200 μL 50% dextran sulfate, 10 μL of yeast tRNA (100 mg mL^-1^), 20 μL 50× Denhardt’s Solution (Thermo Fisher Scientific) and 70 μL dH_2_O. After washing, unbound RNA was digested for 30 min at 37°C using RNase A (20 μg mL^-1^). Immunological detection using DIG antibodies (1:3000 in blocking solution) coupled with alkaline phosphatase and staining procedure with 4-Nitro blue tetrazolium chloride (NBT) and 5-Bromo-4-chloro-3-indolyl-phosphate (BCIP) was done for 24-36 h at room temperature in dark. Hybridization signal analysis was performed using a light microscope BX60 (Olympus).

### Identification of chromatin genes in barley

Identification was done using a subset of 47 *A. thaliana* genes encoding for histones and 12 encoding PRC2 complex subunits. Homology searches were performed using BLAST+ (*73*). The resulting hits were confirmed with reciprocal homology searching using the whole genome of 48,359 *A. thaliana* genes. The candidates were further filtered by standard BLAST+ E-value (≤0.01) and additional parameters counting with the comparison of (1) barley and *A. thaliana* gene length (≥ 40%), and (2) alignment coverage of the hit (≥ 40%).

### Immunostaining and fluorescence *in situ* hybridization (ImmunoFISH), microscopy, and image analysis.x

ImmunoFISH was performed on flow-sorted nuclei from 24 DAP embryos and 8 and 24 DAP endosperm as described (*35*). Preparations were incubated with the rabbit anti-H3K27me3 primary antibody (1:200; Abcam, 195477) at 4°C overnight and the secondary goat anti-rabbit-Alexa Fluor 488 antibody (1:300, Molecular Probes, A11008) at 37°C for 90 min. Barley centromeres were detected with a synthetic 28-mer oligonucleotide (5’-AGGGAGA-3’)_4_ *CEREBA* probe labeled at the 5’ end with Cy3 (Eurofins). Telomeres were visualized with a synthetic 28-mer oligonucleotide probe (5’-CCCTAAA-3)_4_ labeled at the 5 end with Cy5 (Eurofins).

The images were acquired with an AxioImager Z2 microscope (Zeiss, Oberkochen, Germany) equipped with a pE-4000 LED illuminator light source (CoolLED), laser-free confocal spinning disk device (DSD2, Andor, Belfast, UK) and with ×100/1.4 NA Oil M27 Plan-Apochromat (Zeiss) objective. Image stacks of 40-80 slides depending from the C-value of the nucleus, on average, with 0.2 μm z-step were taken separately for each fluorochrome using the appropriate excitation (DAPI λ = 390/40 nm, GFP λ = 482/18 nm, RFP λ = 561/14 nm, Cy5 = 640/14 nm) and emission (DAPI λ = 452/45 nm, GFP λ = 525/45 nm, RFP λ = 609/54 nm, Cy5 = 676/29 nm) filters. The 4.2 MPx sCMOS camera (Zyla 4.2) and the iQ 3.6.1 acquisition software (both Andor) were used to drive the microscope and for fluorescence detection. The images were saved as maximum intensity projection files with Imaris File Converter 9.2.1 (Bitplane, Zurich, Switzerland). Further, Imaris 9.7 functions ‘Surface’ and ‘Spots’ were used for the nucleus surface, immunosignals, and centromere and telomere 3D modeling. Fluorescence intensity measurements were performed in FIJI using the ‘Interactive 3D Surface plot’ plugin.

### Analysis of imprinted genes

The lists of cereal imprinted genes were extracted from published works (*41*–*45*) and their overlaps were visualized using Venn diagrams in R package eulerr (*60*). *H. vulgare* subsp. *spontaneum* accession HOR 12560 was grown as described (*74*) and synchronized for flowering with cv. Morex. The strains were reciprocally crossed, 8 DAP endosperm was manually dissected as described (*52*), total RNA was isolated using Spectrum™ Plant Total RNA Kit (Sigma-Aldrich), and reverse transcribed into cDNA using RevertAid H Minus First Strand cDNA Synthesis Kit (Thermo Fisher Scientific). Primers were designed to amplify 200 – 1,100 bp fragments of the candidate genes harboring informative SNP(s) (table S18) using a standard PCR. The amplicons were gel purified using GeneJet Gel Extraction Kit (Thermo Fisher Scientific) and subjected to Sanger sequencing followed by *in silico* analysis using SnapGene v6.2 software (GSL Biotech LLC).

### Chromatin immunoprecipitation (ChIP)-sequencing and data analysis

Approximately 2 g of 16 DAP endosperm tissue from cv. Morex were isolated in three biological replicates and cross-linked under vacuum in 1% (w/v) formaldehyde for 15 min at room temperature. The cross-linking was quenched by adding glycine to a final concentration of 0.1 M, and the vacuum was applied for 5 min. Endosperm tissue was rinsed with water and frozen in liquid nitrogen. ChIP was performed as described previously (*75*), with the following modifications. Briefly, isolated nuclei were resuspended in Nuclei lysis buffer (50 mM Tris-HCl pH 8.0, 1 mM EDTA, 0.5% SDS, cOmplete™, EDTA-free Protease Inhibitor Cocktail (Roche)) and incubated at 4°C under gentle agitation for 20 min. The chromatin solution was sonicated using a Biorupter Plus (Diagenode) with 10 cycles of 30 sec pulse/90 sec cooling at high power at 4°C. The resulting sheared chromatin was pooled and diluted 1:4 with ChIP dilution buffer (16.7 mM Tris-HCl pH 8.0, 167 mM NaCl, 1.25 mM EDTA, 1.25% Triton X-100, 1× cOmplete™, EDTA-free Protease Inhibitor Cocktail). 600 μl aliquots of diluted chromatin were incubated with 7 μl of the rabbit anti-trimethyl-Histone H3 (Lys27) antibody (Millipore, 07-449) in a rotator at 4°C overnight. Samples without antibody were used as negative controls. The next day, 40 μl of the Dynabeads Protein A (Invitrogen) were added to each tube, and the samples were further incubated for 2 hours. Afterward, beads were washed with a sequence of buffers, and immune complexes were eluted as described. Control chromatin aliquots (‘input DNA’) were taken prior to immunoprecipitation. Reverse crosslinking was performed for all samples, and DNA was extracted and purified using the ChIP DNA Clean & Concentrator™ kit (Zymo Research). Sequencing libraries were prepared and 150 bp pair-end reads were sequenced using Illumina NovaSeq 6000 platform (Illumina) by Novogene.

The raw reads of samples sequenced in our study and also publicly available H3K27me3 ChIP-seq data from shoot (*36*) were trimmed for adaptors and aligned to the Morex reference genome, duplicates were removed using MarkDuplicates in GATK (*76*) and the peak calling was performed using MACS2 (*77*). The peaks were filtered (fold enrichment ≥ 5) and tested for differential signal intensity between endosperm and shoot samples using R package MAnorm2 (*78*). Testing was performed in genomic intervals of size 2000 nt and intervals with differential signal intensities localized in coding regions or 2000 bp upstream were related to genes. GO term enrichment analysis was performed by web based tool g:Profiler (*79*) using barley GO annotation for Morex V3 available at Enembl Plants.

## Accession numbers

Unique identifiers for all genes and their products mentioned in the text are provided in table S19.

## Acknowledgments

We thank H. Tvardíková for technical assistance, Z. Bursová for plant care, and P. Navrátil for IT support. Computational resources were supplied by the project “e-Infrastruktura CZ” (e-INFRA CZ LM2018140) supported by the Ministry of Education, Youth and Sports of the Czech Republic.

## Author contributions

M.Ko. performed sample preparation and RNA extraction, data analysis and interpretation. M.Ko. and I.V. performed RNA *in situ* hybridization. A.N. performed immunostaining and fluorescence *in situ* hybridization, microscopy, image analysis and barley crosses. J.Z. performed identification of chromatin genes. B.S. performed ChIP and contributed to data analysis. N.J.P., E.E. and A.Pa. prepared a seed view in barley ePlant. K.K. and M.Kr. performed validation of imprinted genes. R.S. and J.Š. supervised parts of the project. A.Pe. conceived, designed, and supervised the project. A.Pe and M.Ko. wrote the manuscript. All authors have seen and agreed upon the final version of the manuscript.

## Data and materials availability

All original RNA-seq and ChIP-seq data from this article can be found in the Gene Expression Omnibus under accession numbers GSE233316 and GSE238237, respectively.

