## Supplementary Materials for "The transcriptome landscape of developing barley seeds reveals H3K27me3 dynamics in endosperm tissues"

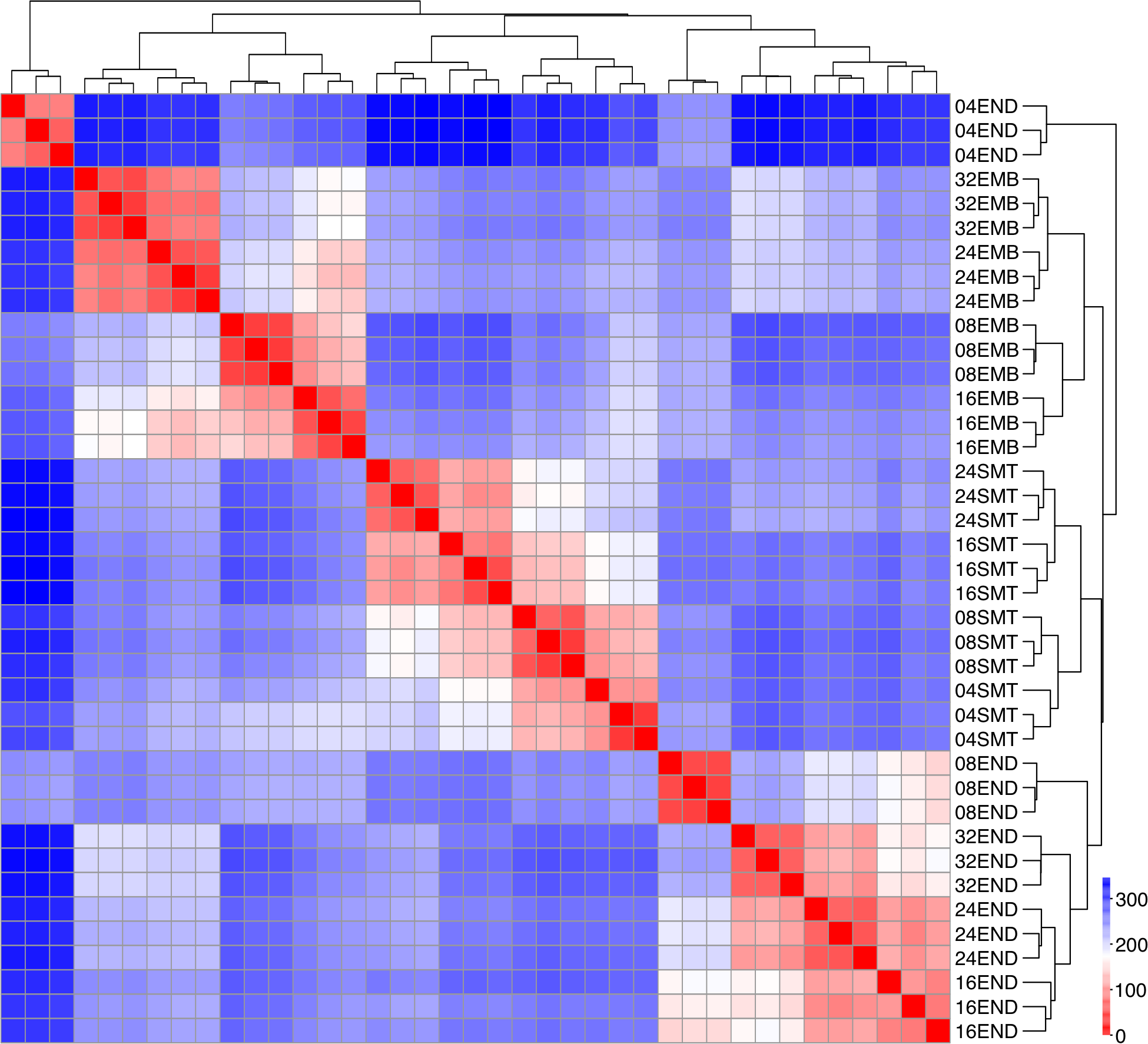

Fig. S1.

Sample distance heatmap of 39 transcriptomic samples from three seed tissues hierarchically clustered based on their similarity. Biological replicates are shown on the right side of heatmap. The scale shows sample distance.

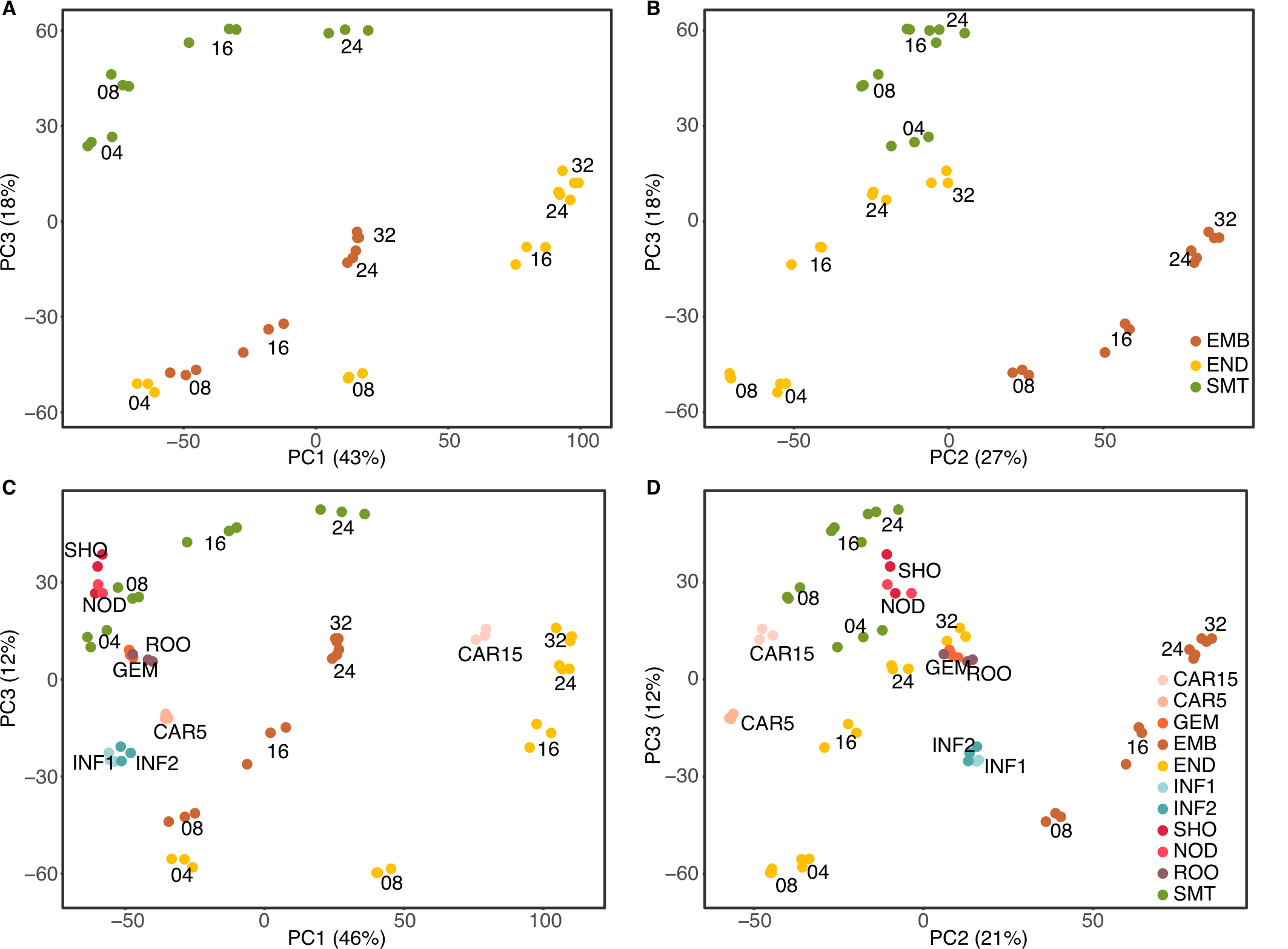

Fig. S2.

Principal component (PC) analysis of seed transcriptomes. (**A-B**) Combinations with PC3 explaining in total 88% of the variability. (**C-D**) Comparison with 8 tissue-specific transcriptomes of vegetative tissues done by IBGSC, 2012. Combinations with PC3 explaining 79% of the variability. Embryo (orange), endosperm (yellow) and seed maternal tissues (green). The numbers within the graph indicate DAP and three close spots represent biological triplicates. ROO – root, GEM – germinating embryo, NOD – nodule, SHO – shoot, INF1 and INF2 – developing inflorescence of 5 and 10 mm length, CAR5 and CAR15 – caryopsis 5 and 15 DAP. The numbers next to PCs indicate variance.

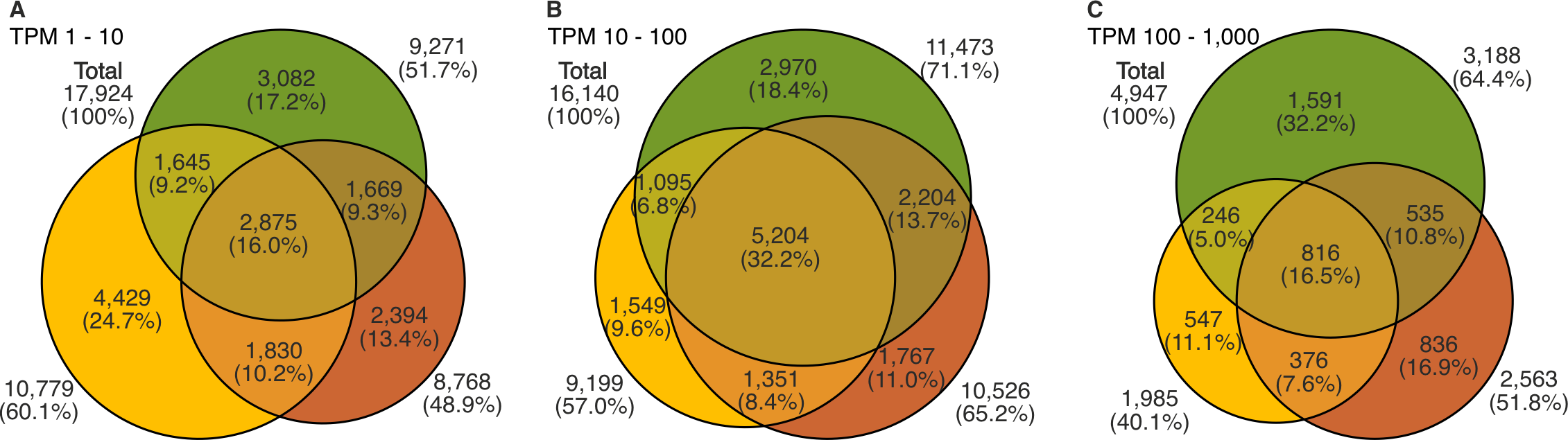

Fig. S3.

Venn diagram showing numbers of tissue-specific and shared genes among seed tissues expressed at different level (**A**) 1-10 TPM. (**B**) 10-100 TPM (**C**) 100-1,000 TPM. Embryo (orange), endosperm (yellow), seed maternal tissues (green). TPM – transcripts per million.

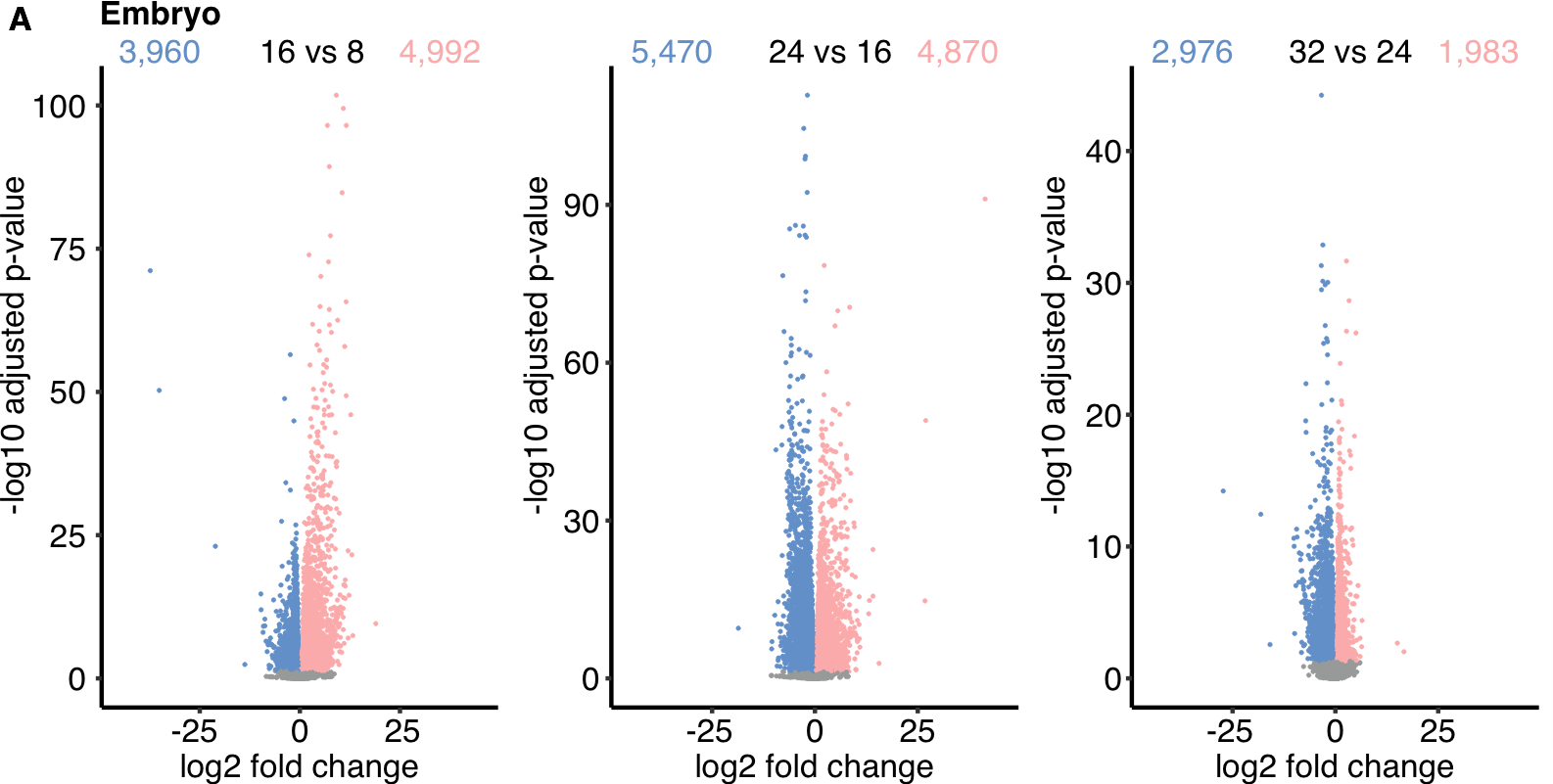

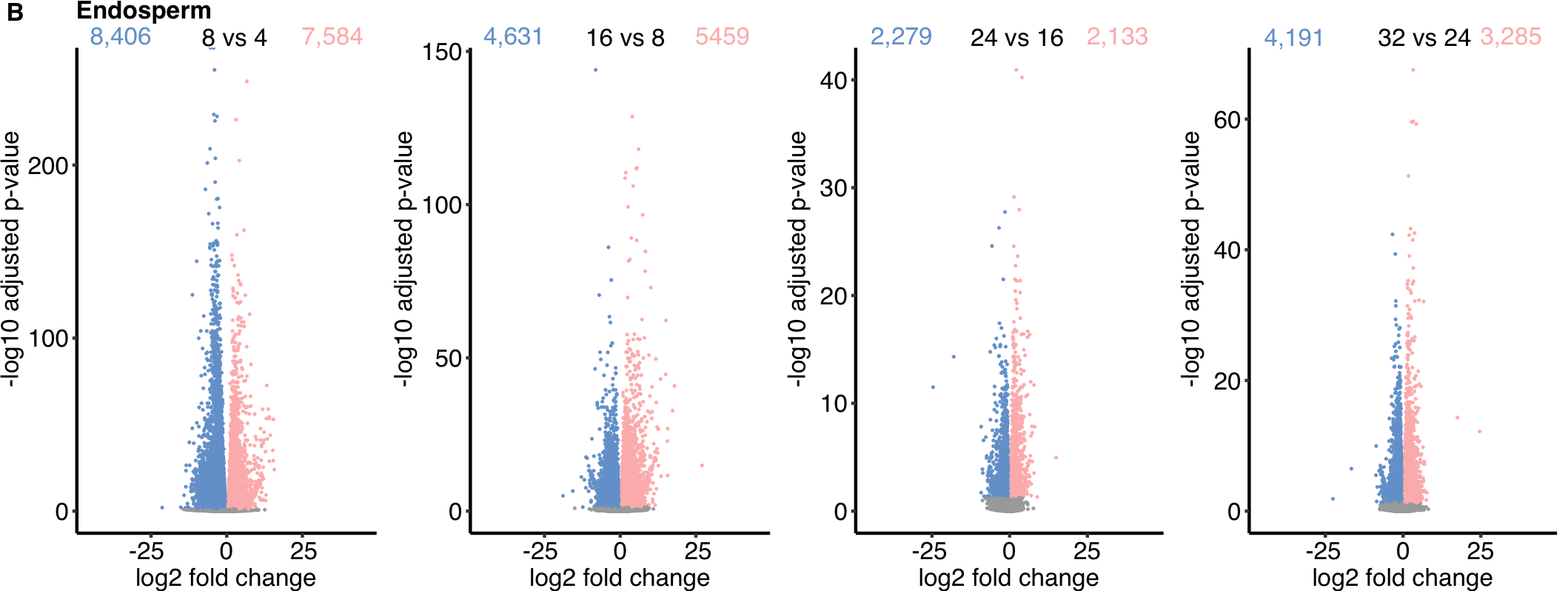

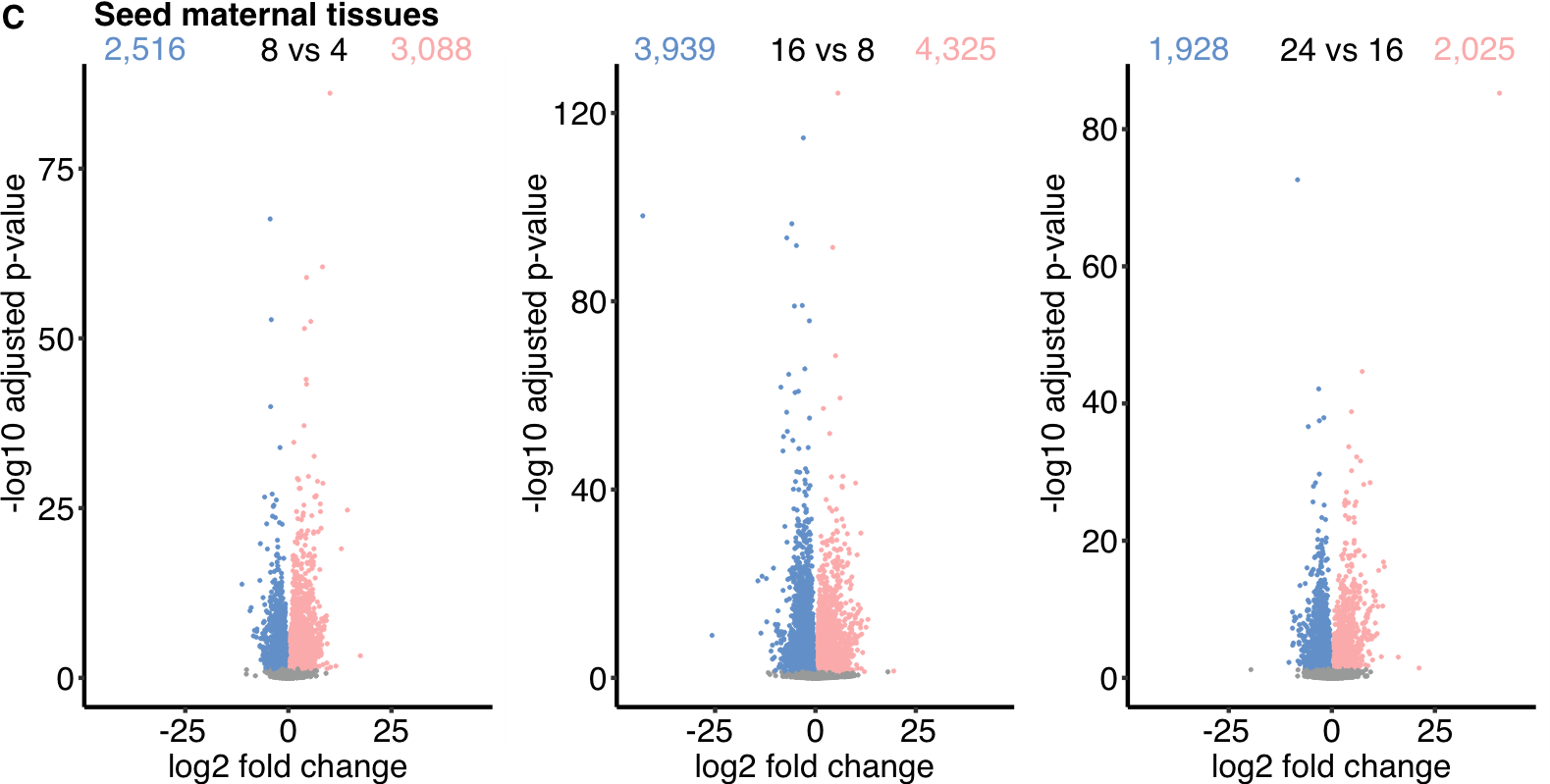

Fig. S4.

Volcano plots showing differentially expressed genes (DEGs) in consecutive developmental transitions in (**A**) embryo, (**B**) endosperm, and (**C**) seed maternal tissues. Genes passing the threshold (padj < 0.05) are colored in blue (down-regulated) or red (up-regulated). Grey dots indicate genes with non-significant expression changes.

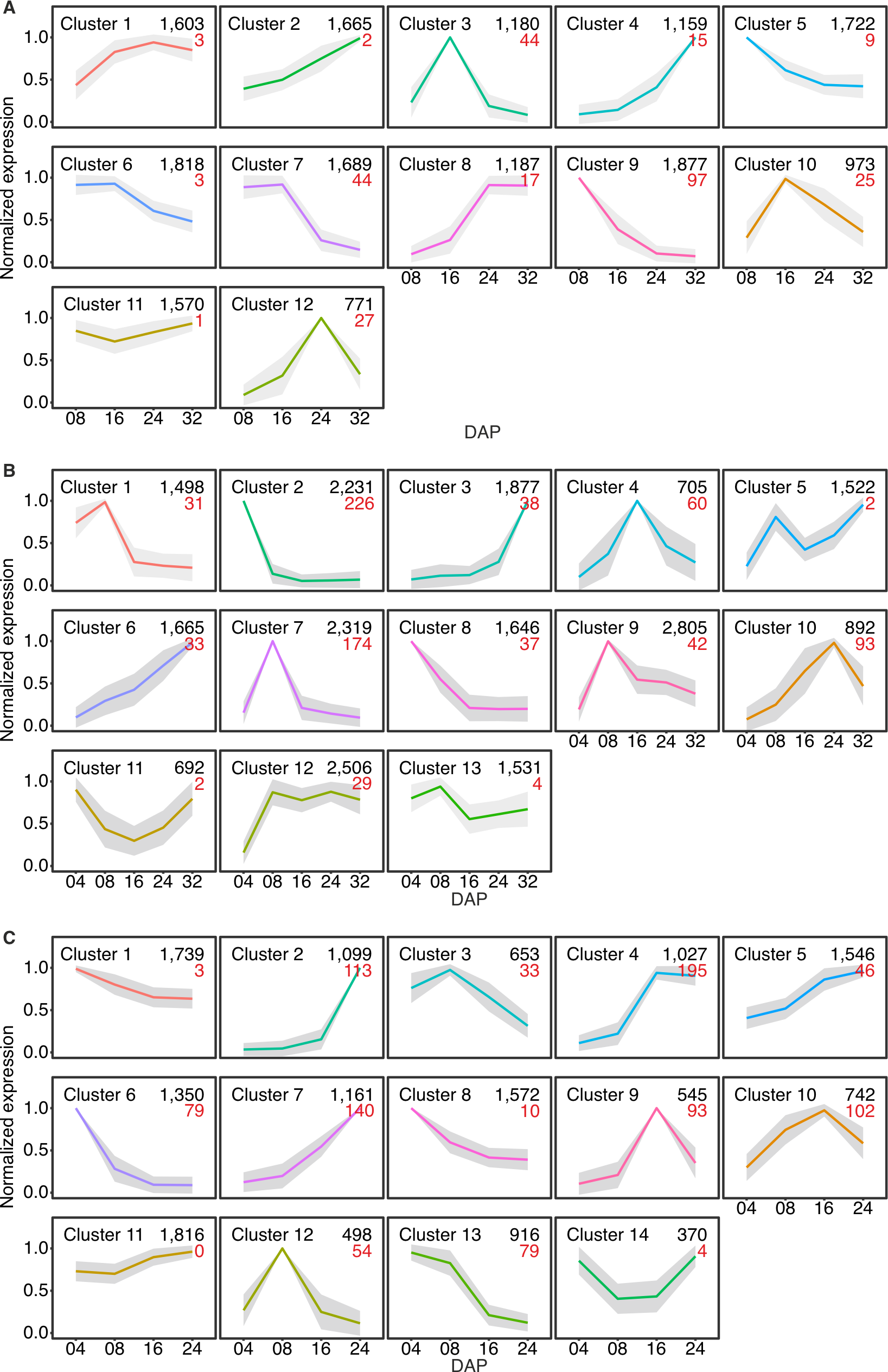

Fig. S5.

K-means co-expression clusters of DEGs, identified in (**A**) embryo, (**B**) endosperm, and (**C**) seed maternal tissues. The black numbers in the upper right corner indicate gene count in individual clusters and the red numbers show the count of tissues-specific genes relative to other seed tissues used in this study. DAP – days after pollination.

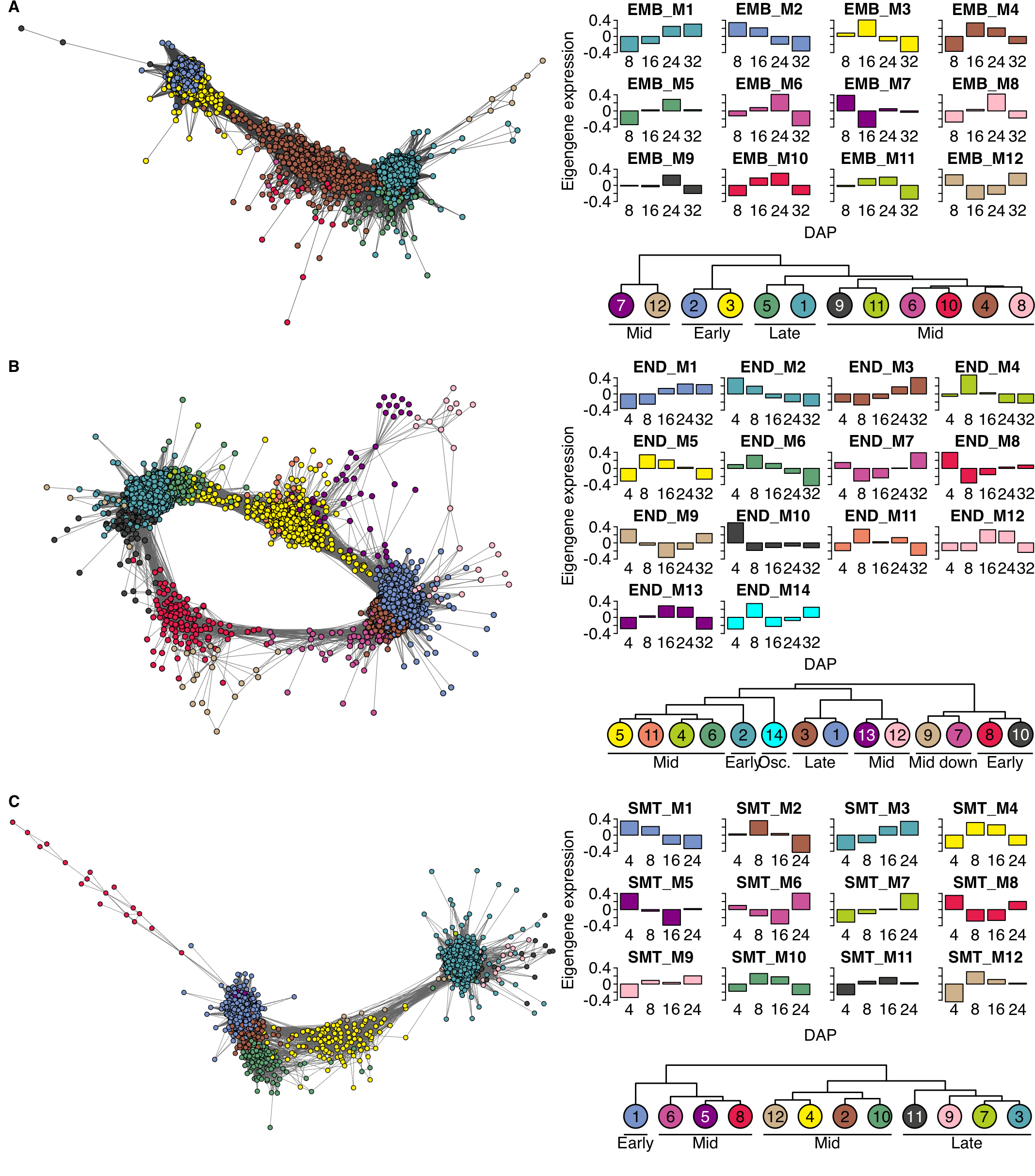

Fig. S6.

Display of weighted gene co-expression network analysis, corresponding modules and its hierarchical clustering in embryo (EMB) (**A**), endosperm (END) (**B**) and seed maternal tissues (SMTs) (**C**). The network shows individual genes (points) and expression similarities (lines) passing criteria (weight > 0.2). Individual modules are marked with different colors and connected in temporal manner. Bar plots show eigengene expression in each module. Individual modules are clustered based on the eigengene expression and divided into early, mid, late, mid down and oscillating. Colors and numbers in the network, bar plots and hierarchical clusters correspond to each other. DAP – days after pollination. Osc. – oscillating.

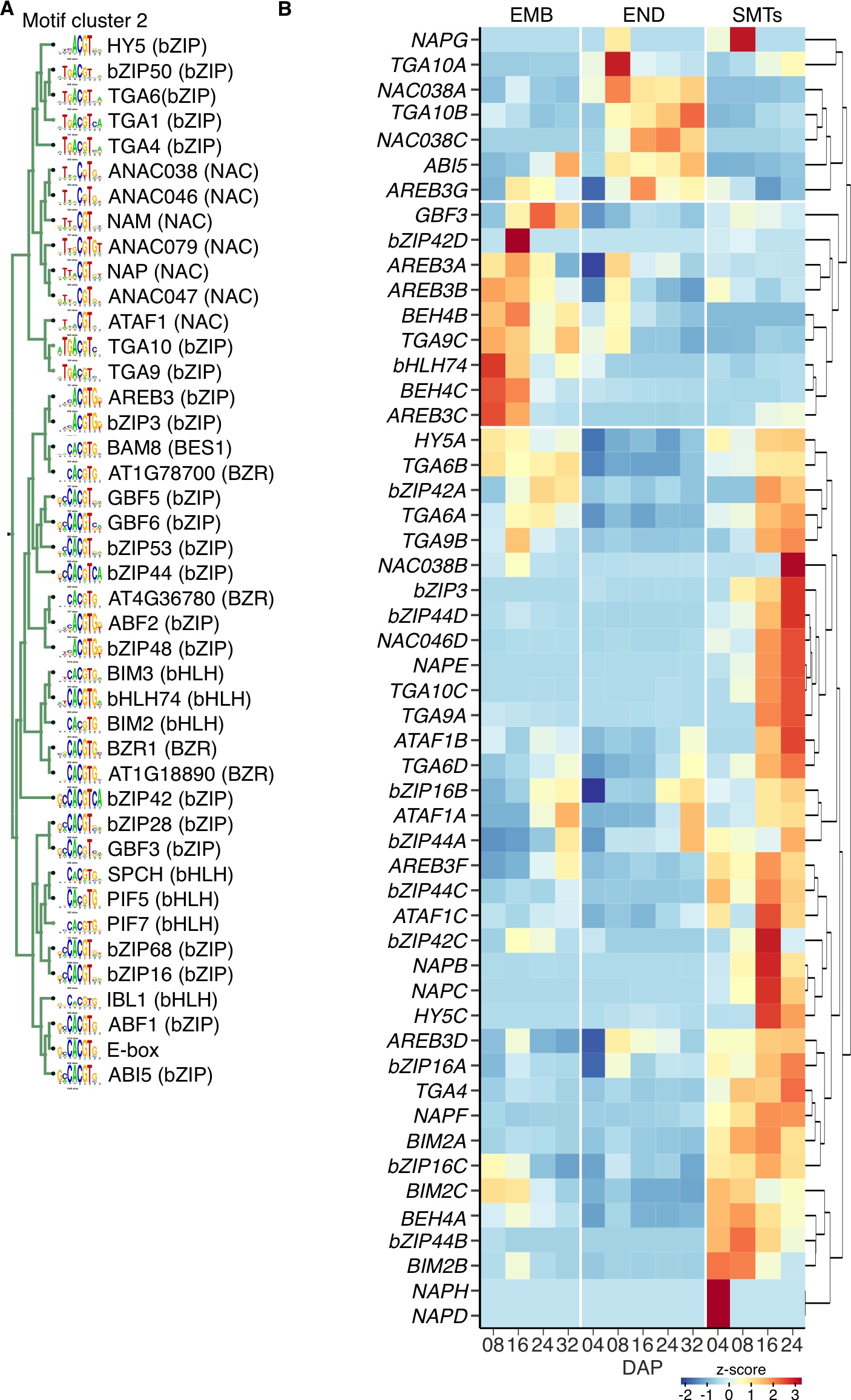

Fig. S7

(**A**) Detailed clustering of TF binding sequence motifs in MC9 as to their similarity. The motifs of Arabidopsis TFs and their families (in parentheses) are shown. (**B**) Heatmap of hierarchically clustered expression for barley orthologs of Arabidopsis TFs from (A) in embryo (EMB), endosperm (END) and seed maternal tissues (SMTs; source data in table S7).

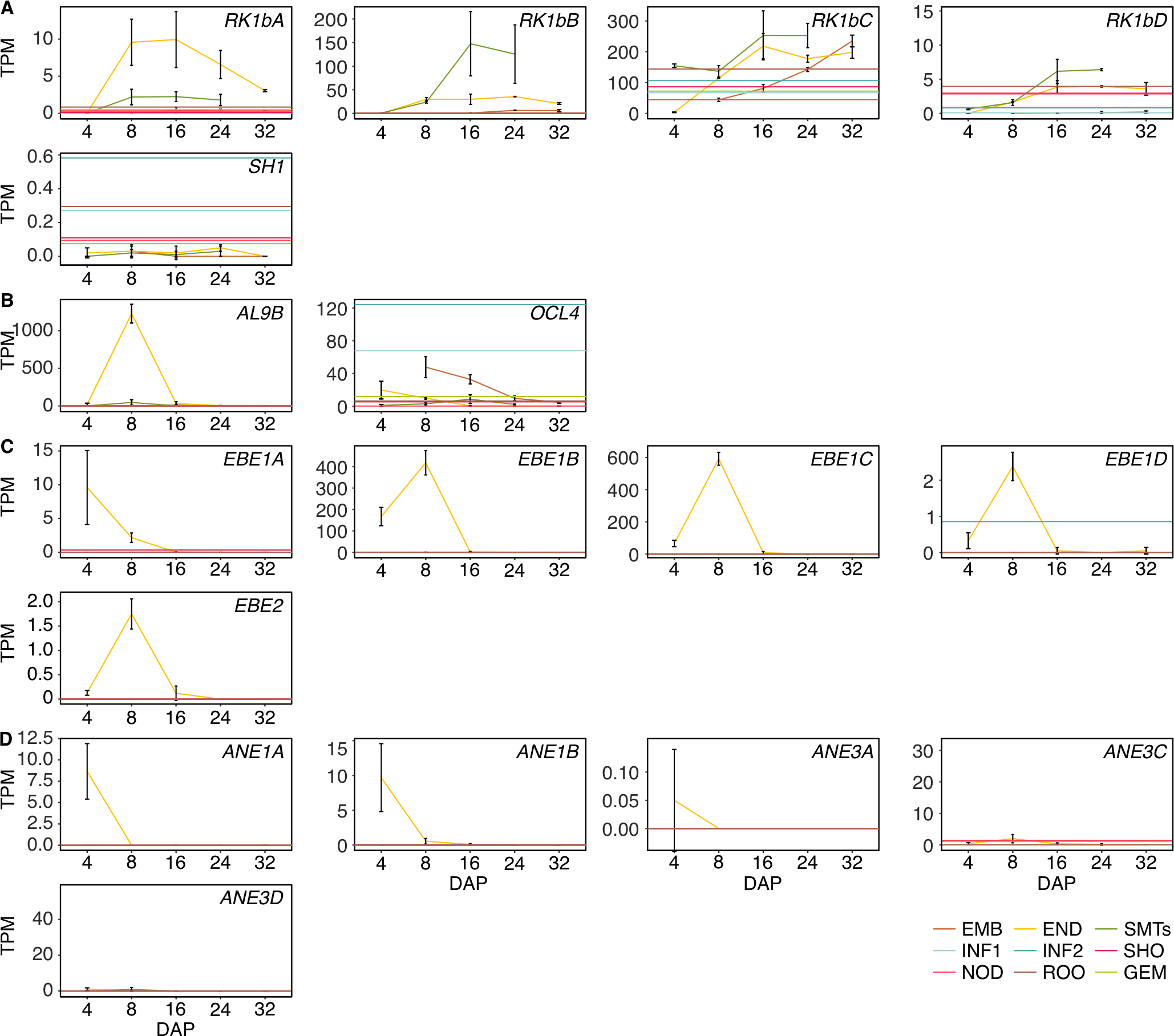

Fig. S8

Expression profiles of barley orthologs of selected endosperm marker genes in other cereals grouped according to the domain of expression (**A**) central starchy endosperm, (**B**) aleurone layer, (**C**) basal endosperm transfer layer and (**D**) embryo surrounding region across different barley tissues. EMB – embryo, END – endosperm, SMTs – seed maternal tissues, ROO – root, GEM – germinating embryo, NOD – nodule, SHO – shoot, INF1 and INF2 – developing inflorescence 5 and 10 mm, CAR5 and CAR15 – caryopsis 5 and 15 DAP. Error bars indicate standard deviation.

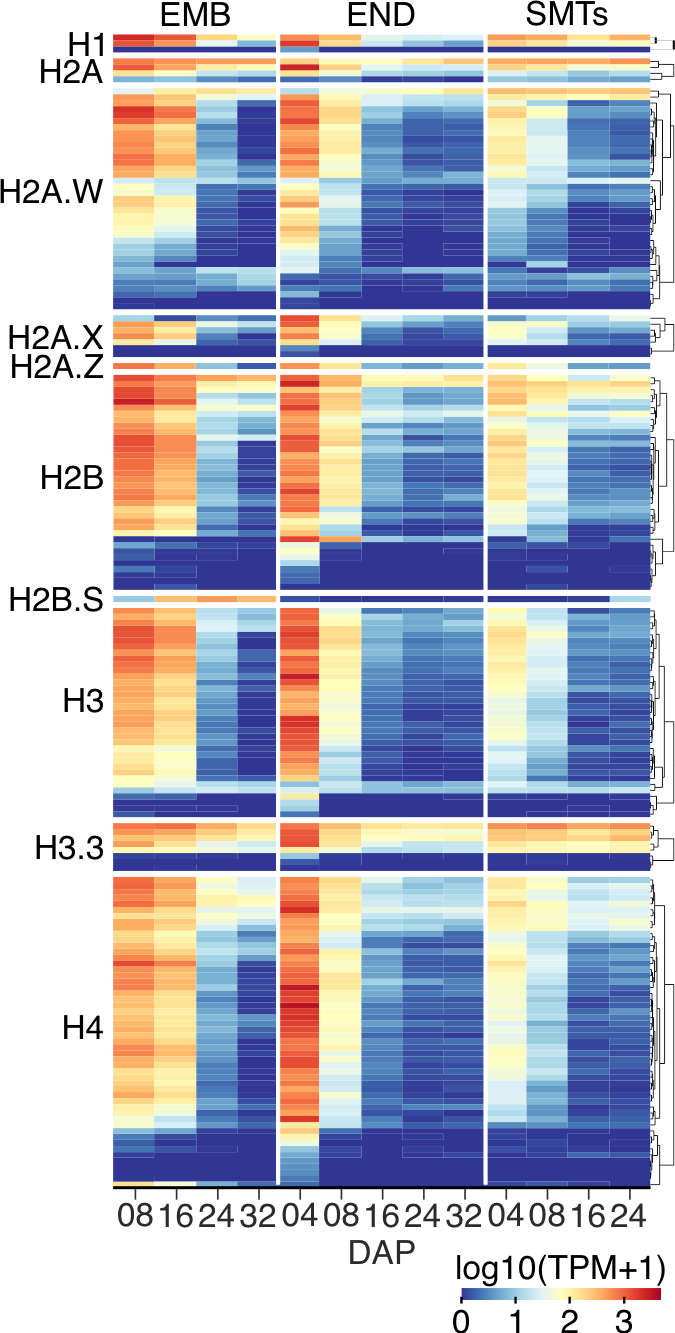
Fig. S9

Heatmap of expression from 175 histone genes in the embryo (EMB), endosperm (END), and seed maternal tissues (SMT) at different days after pollination (DAP). The heatmap is partitioned into groups according to histone variants and hierarchically clustered within the groups. Source data are provided in table S10.

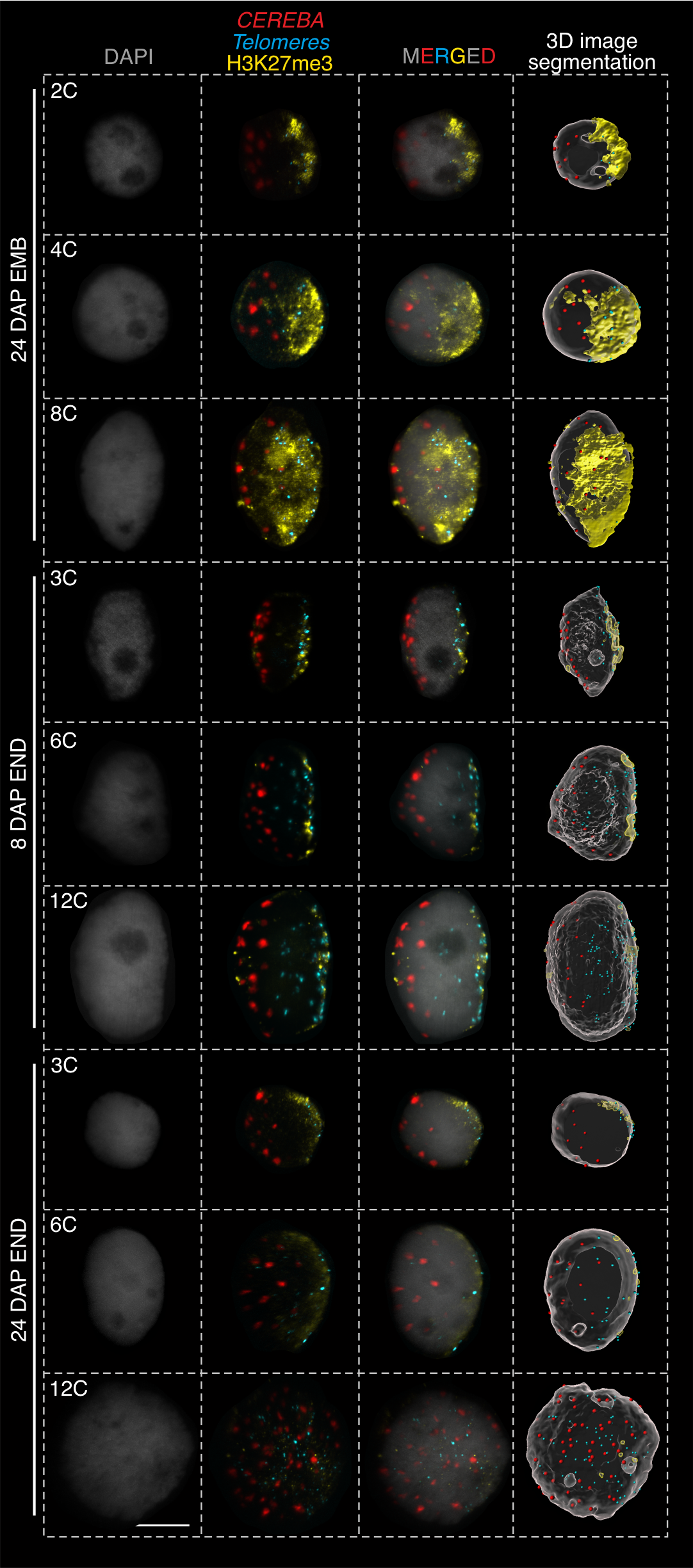
**Fig. S10**

Raw images and 3D image segmentation pictures of embryo and endosperm nuclei of different C-values collected at 8 and 24 DAP after immunostaining with anti-H3K27me3 antibody (yellow), followed by fluorescence *in situ* hybridization for *CEREBA* centromeric (red) and telomeric (blue) repeats. The last column shows 3D image segmentation of the nucleus surface, immunostaining and FISH signals. This allows visualizing the spatial distribution of the H3K27me3 histone modification, centromeres, and telomeres within the nucleus. Scale bar = 10 µm.

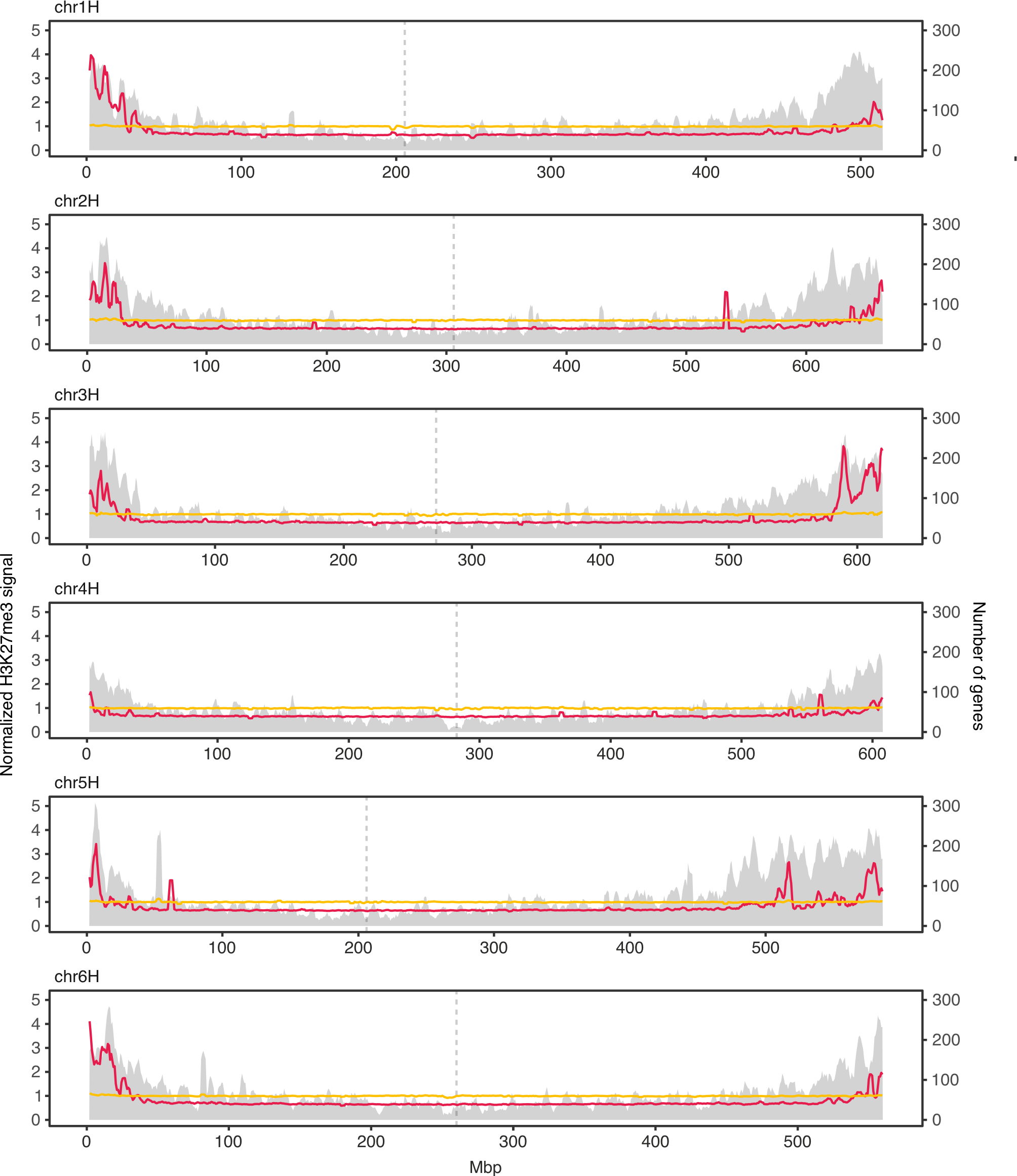

Fig. S11

Sliding window across chromosome 1H-6H showing normalized signal abundance of H3K27me3 in endosperm (yellow) and 10-cm whole seedling (*51*) (magenta) and number of genes (secondary y-axis, gray).

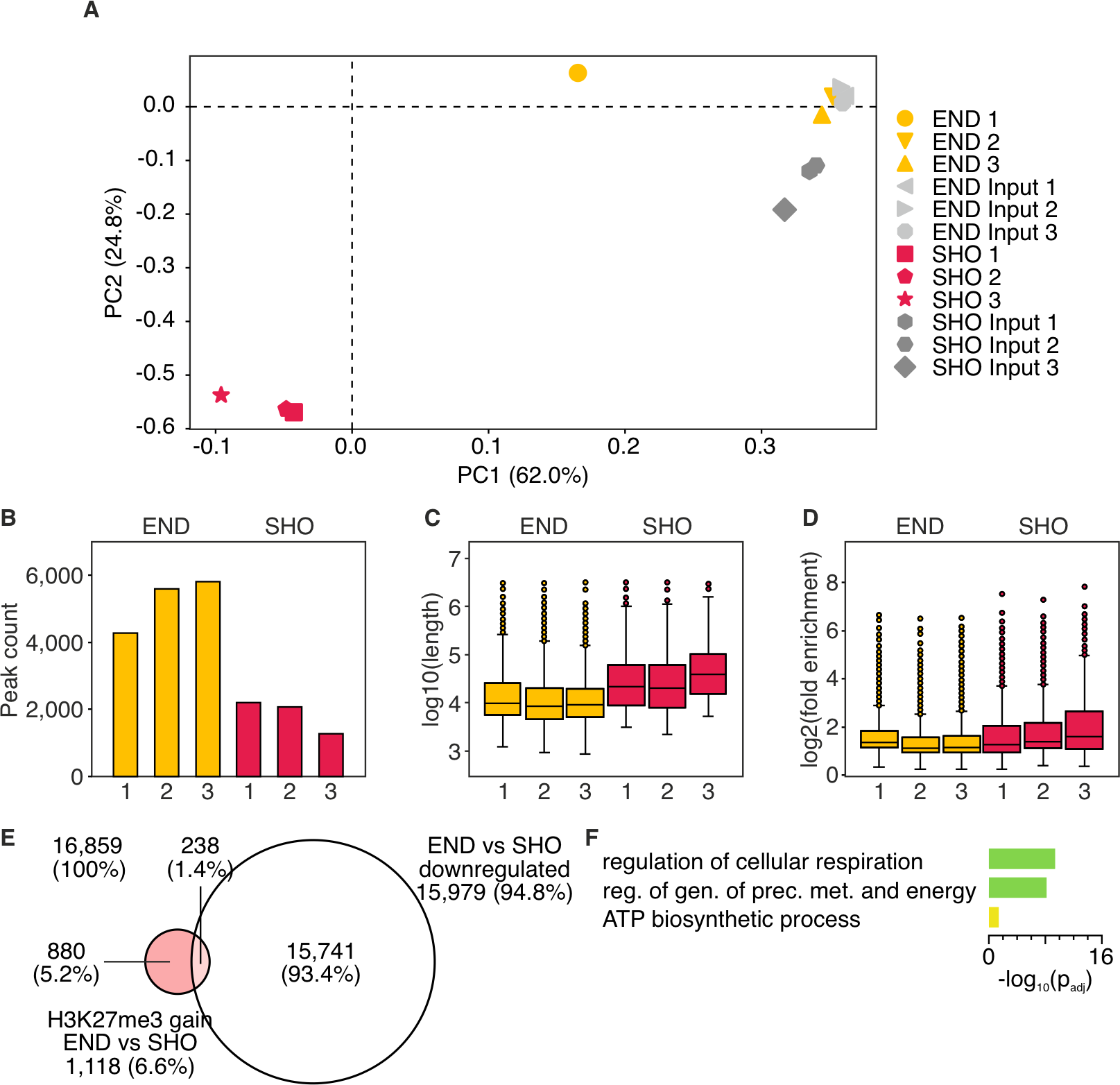

Fig. S12

(**A**) The variance of the ChIP-seq samples used in this study represented by the principal component (PC) analysis. END - 16 DAP endosperm, SHO - 10-cm seedling (*36*). The numbers next to PCs indicate variance. (**B-D**) ChIP-seq peak statistics showing (**B**) the number of called peaks, (**C**) distribution of peak length and (**D**) fold enrichment over input. (**E**) Venn diagram showing the number of genes with gain of H3K27me3 in 10-cm seedlings found by ChIP-seq and genes up-regulated in at least one endosperm sample (8, 16, 24, or 32 DAP) as determined by RNA-seq. (**F**) Gene ontology (GO) term enrichment of genes with loss of H3K27me3 and significant up-regulation in endosperm (source data in table S15).

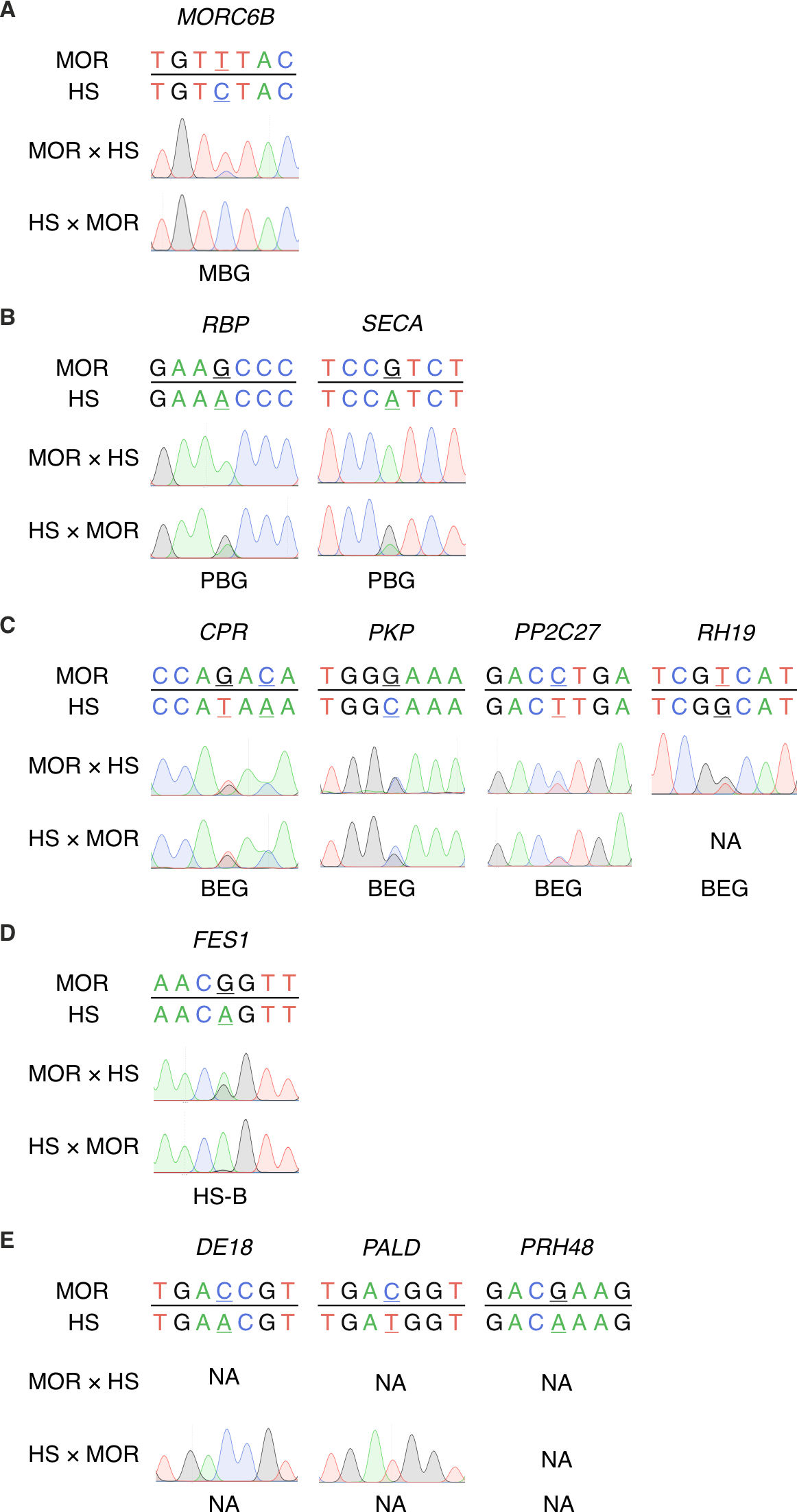

Fig. S13

Overview of genes that were not confirmed as imprinted in based on the analysis of F1 reciprocal crossed of Morex (MOR) × wild barley *H. spontaneum* accession HOR 12560 (HS) in 8 DAP endosperm tissues by Sanger sequencing. (**A**) maternally biased genes, (**B**) paternally biased genes, (**C**) biparentally expressed genes, (**D**) *H. spontaneum* biased and (**E**) information not available. The informative SNPs are underlined.

Table S1. (separate file)

Gene ontology (GO) term category enrichment based on the subset of genes with TPM > 1,000 in endosperm tissues.

Table S2. (separate file)

Gene ontology (GO) term enrichment in for down-regulated DEGs (FDR adjusted p-value <0.05) at subsequent experimental points in embryo (EMB), endosperm (END) and seed maternal tissues (SMTs). NS - not significant. Annotated – number of genes in the genome having specific GO term. Expected – number of genes expected to have specific GO term in the subset. Significant – number of genes having a specific GO term in the subset.

Table S3. (separate file)

Gene ontology (GO) term enrichment in for up-regulated DEGs (FDR adjusted p-value <0.05) at subsequent experimental points in embryo (EMB), endosperm (END) and seed maternal tissues (SMTs). NS - not significant. Annotated – number of genes in the genome having specific GO term. Expected – number of genes expected to have specific GO term in the subset. Significant – number of genes having a specific GO term in the subset.

Table S4. (separate file)

List of motifs assigned to different motif clusters.

Table S5. (separate file)

Contribution of each motif collection to the motif clusters identified by RSAT.

Table S6. (separate file)

List of barley genes found by homology search using Arabidopsis proteins with binding motifs assigned to motif cluster 2. pident - percentage of identical positions. length – alignment length (sequence overlap). mismatch – number of mismatches. gapopen – number of gap openings. qstart – start of the alignment in query. qend – end of the alignment in query. sstart – start of the alignment in subject. send – end of the alignment in subject. evalue – expect value. bitscore – bit score.

Table S7. (separate file)

List of barley genes found by homology search using Arabidopsis proteins with binding motifs assigned to motif cluster 9. pident - percentage of identical positions. length – alignment length (sequence overlap). mismatch – number of mismatches. gapopen – number of gap openings. qstart – start of the alignment in query. qend – end of the alignment in query. sstart – start of the alignment in subject. send – end of the alignment in subject. evalue – expect value. bitscore – bit score.

Table S8. (separate file)
List of endosperm domains markers in barley found by homology search using known rice and maize proteins. pident - percentage of identical positions. length – alignment length (sequence overlap). mismatch – number of mismatches. gapopen – number of gap openings. qstart – start of the alignment in query. qend – end of the alignment in query. sstart – start of the alignment in subject. send – end of the alignment in subject. evalue – expect value. bitscore – bit score.

Table S9. (separate file)

List of barley histone and PRC2 genes found by homology search using Arabidopsis proteins. pident - percentage of identical positions. length – alignment length (sequence overlap). mismatch – number of mismatches. gapopen – number of gap openings. qstart – start of the alignment in query. qend – end of the alignment in query. sstart – start of the alignment in subject. send – end of the alignment in subject. evalue – expect value. bitscore – bit score.

Table S10. (separate file)

Seed transcriptomic data for histone genes in barley.

Table S11. (separate file)

Seed transcriptomic data for core PRC2 genes in barley.

Table S12. (separate file)

List of genes significantly up-regulated in RNA-seq and with significantly reduced H3K27me3 in ChIP-seq in 16 days after pollination endosperm (END) compared to shoot (SHO).

Table S13. (separate file)
Gene ontology (GO) term enrichment for genes up-regulated in endosperm RNA-seq and with significantly reduced H3K27me3 in ChIP-seq in 16 days after pollination endosperm (END) compared to shoot (SHO). Source – category of GO terms. Term size – number of genes in the genome having specific GO term. Query size – number of analyzed genes in specific GO term category. Intersection size – number of analyzed genes having a specific GO term in the subset.

Table S14. (separate file)
List of genes significantly down-regulated in RNA-seq and with significantly increased H3K27me3 in ChIP-seq in 16 days after pollination endosperm (END) compared to shoot (SHO).

Table S15. (separate file)
Gene ontology (GO) term enrichment for genes down-regulated in endosperm RNA-seq and with significantly increased H3K27me3 in ChIP-seq in 16 days after pollination endosperm (END) compared to shoot (SHO). Source – category of GO terms. Term size – number of genes in the genome having specific GO term. Query size – number of analyzed genes in specific GO term category. Intersection size – number of analyzed genes having a specific GO term in the subset.

Table S16. (separate file)

List of barley candidate imprinted genes found by homology search using known maize, rice, and wheat imprinted genes. pident - percentage of identical positions. length – alignment length (sequence overlap). mismatch – number of mismatches. gapopen – number of gap openings. qstart – start of the alignment in query. qend – end of the alignment in query. sstart – start of the alignment in subject. send – end of the alignment in subject. evalue – expect value. bitscore – bit score.

Table S17. (separate file)

Seed transcriptomic data for barley homologs of imprinted genes identified in at least two other cereals.

Table S18. (separate file)

List of primers used in this study.

Table S19. (separate file)

List of genes and identifiers (IDs) used in the text.

Data S1. (separate file)

RNA-seq raw read counts for embryo (EMB), endosperm (END) and seed maternal tissues (SMTs).

Data S2. (separate file)

Expression changes between individual subsequent days after pollination in embryo (EMB), endosperm (END) and seed maternal tissues (SMTs). NA - not available.

Data S3. (separate file)

Assignment of differentially expressed genes from embryo (EMB), endosperm (END) and seed maternal tissues (SMTs) to individual k-means clusters (K). NA - not available.

Data S4. (separate file)

Gene assignment to WGCNA modules in embryo (EMB), endosperm (END) and seed maternal tissues (SMTs). (NA - not available).

Data S5. (separate file)

Identified plant motifs within WGCNA modules in embryo, endosperm and seed maternal tissues.

Data S6. (separate file)

List of genomic intervals with significantly differential signal intensity of H3K27me3 in endosperm (END) compared to shoot (SHO) in ChIP-seq. SHO.mean, END.mean – mean signal intensities of shoot and endosperm, respectively. Mval – difference in mean signal intensity (log2 foldChange). Mval.se – standard error associated with the Mval. Mval.t – the ratio of Mval to Mval.se. pval – two sided p-value for the statistical significance. padj – p-value adjusted for multiple testing with the "BH" method.

Data S7. (separate file)

List of genes with significantly differential signal intensity of H3K27me3 in endosperm (END) compared to shoot (SHO).

Data S8. (separate file)

List of transcription factors and their assignments to families.
